# Mouse helpers ensure maternal-infant survival

**DOI:** 10.1101/2022.12.26.521927

**Authors:** Luisa Schuster, Violet J. Ivan, Diana M. Gilly Suarez, Renee Henderson, Asha Caslin, Jessica L. Minder, Gurket Kaur, Shreya Sankar, Deepasri Ananth, Matilda Kirk, Sarah B. Winokur, Latika Khatri, Paola Leone, Karen E. Adolph, Robert C. Froemke, Adam Mar

## Abstract

Parental care is required for offspring survival, because infants require nearly-continual oversight for extensive periods. Parents must balance caretaking with their own survival, and benefit from help provided by other experienced adults. We built a system for 24/7 long-term monitoring of wild-type or oxytocin receptor knockout (OXTR-KO) mouse mothers (‘dams’) over litters. Some wild-type dams had high litter survival rates, but others consistently lost pups due to neglect and hypothermia. Maternal caretaking in low-pup-survival dams improved after co-housing with an experienced dam and litter, from reorganized maternal behavior including increased nest-building. In contrast, singly-housed OXTR-KOs died in childbirth and pups died due to prolonged parturition. Co-housing with a lactating female prevented OXTR-KO maternal-infant mortality, because the other female acted as a ‘midwife’ by removing and cleaning pups from the pregnant dam. Thus for single mothers that continually lose litters or die in labor, maternal-infant survival is enhanced by experienced helpers.

## Introduction

Parental behavior is essential for the well-being of offspring and the survival of the species^1–2^. Adult animals can become highly sensitive to the needs of their infants to adequately care for them, as infants require intense continuous protection for days to years depending on species. Mouse pups, for example, rely on their mother for sustenance, thermoregulation, and protection, supported by successful nest-building^3–5^. However, pup mortality might result if dams are neglectful or provide inadequate care during the postnatal period. Parental animals must balance caregiving behaviors with other behaviors required for their own survival, making single parenting especially challenging. Previous studies of rodents and other species report a remarkably high degree of infant mortality, both outdoors in native habitats and animals housed in laboratory conditions-although it remains controversial how much infant mortality occurs for isolate vs communally-rearing mothers, and to what degree this is genetically predisposed, influenced by the environment, or shaped by life experiences^6–10^.

The hormone oxytocin is important for coordinating and regulating pro-social behavior and maternal care. Oxytocin is a nine amino-acid neuropeptide synthesized mainly in the hypothalamus, with a single G-protein-coupled receptor in the mammalian genome^11–14^. Oxytocin neurons are activated by social interactions such as somatic contact^15^, visual observation of experienced maternal animals^16^, and the sounds of infant vocalizations^17,18^, with release of oxytocin into different brain regions hypothesized to promote behaviors such as maternal care and co-parenting by increasing the salience of social cues^19^. Oxytocin signaling is also critical for a number of physiological processes, including two mechanisms fundamental for mammalian reproduction: milk ejection during nursing and uterine smooth muscle contractions for parturition^12,13,19^. Knocking out the oxytocin receptor in mice or voles greatly reduced litter sizes and pup survival^20,21^, likely due to difficulties feeding neonates in the absence of mammary contractile responses. However, despite the widespread use of oxytocin as a uterotonic for aiding human labor, these studies reported that oxytocin receptor knockout (OXTR-KO) mice and voles seem to have normal parturition^20,21^. It is unknown how these animals lacking oxytocin receptor signaling give birth, especially as receptor knockout mice were shown to lack uterine contractions in response to either oxytocin or vasopressin application^20^. This issue is not unique to rodents; across species, there is a substantial lack of information about the details of labor and birthing mechanisms in most non-human species including mice^22–25^. Lack of data on parturition, mother and infant survival, and the importance (or lack thereof) of oxytocin signaling for birthing and parental care make it challenging to translate findings in non-human species to treatments for improving human mother-infant health and wellness^26,27^.

Advances in hardware and software now make it feasible to collect, store, and analyze datasets on a massive scale. While most behavioral experiments collect data at specific timepoints or assay certain functions, in principle it is now possible to monitor the spontaneous, every-day behaviors of animals continuously over long periods of time. Our goal was to build a low-cost, multi-camera behavioral monitoring system to characterize the family life of lab mice. By studying singly-housed or pair-housed wild-type or OXTR-KO dams, here we aimed to determine how dams give birth, when and how pup mortality occurs (due to parental neglect or other factors), and if mouse mothers might learn from certain experiences to improve maternal care over subsequent litters.

## Results

### A substantial subset of mouse dams repeatedly lost pups due to neglect

We designed a high-resolution system to monitor parental care and family life in mice with multiple cameras, recording from homecages continuously 24 hours/day for weeks to months across consecutive litters (**Fig. 1A; Supplementary Video 1**). Individual cages were contained within custom-designed environmental control boxes, containing up to five cameras that recorded behavior from the top, side, and underneath the nest. We also used infrared cameras to record in darkness and a thermal imaging camera to monitor temperature. Videos were combined with an inexpensive, open-source system to control LED light-strips to provide a naturalistic light/dark cycle, together with temperature and humidity sensors connected to an Arduino circuit board that controls two low-noise fans, and adjusts their speed to maintain the enclosure temperature at 26-28 °C (**Fig. 1B**), a level close to adult mouse thermoneutrality of ∼30°C^5,28^. Data were saved to local servers and streamed remotely for quantitative behavioral analysis based on machine learning and multi-rater human manual annotation.

**Figure 1.**
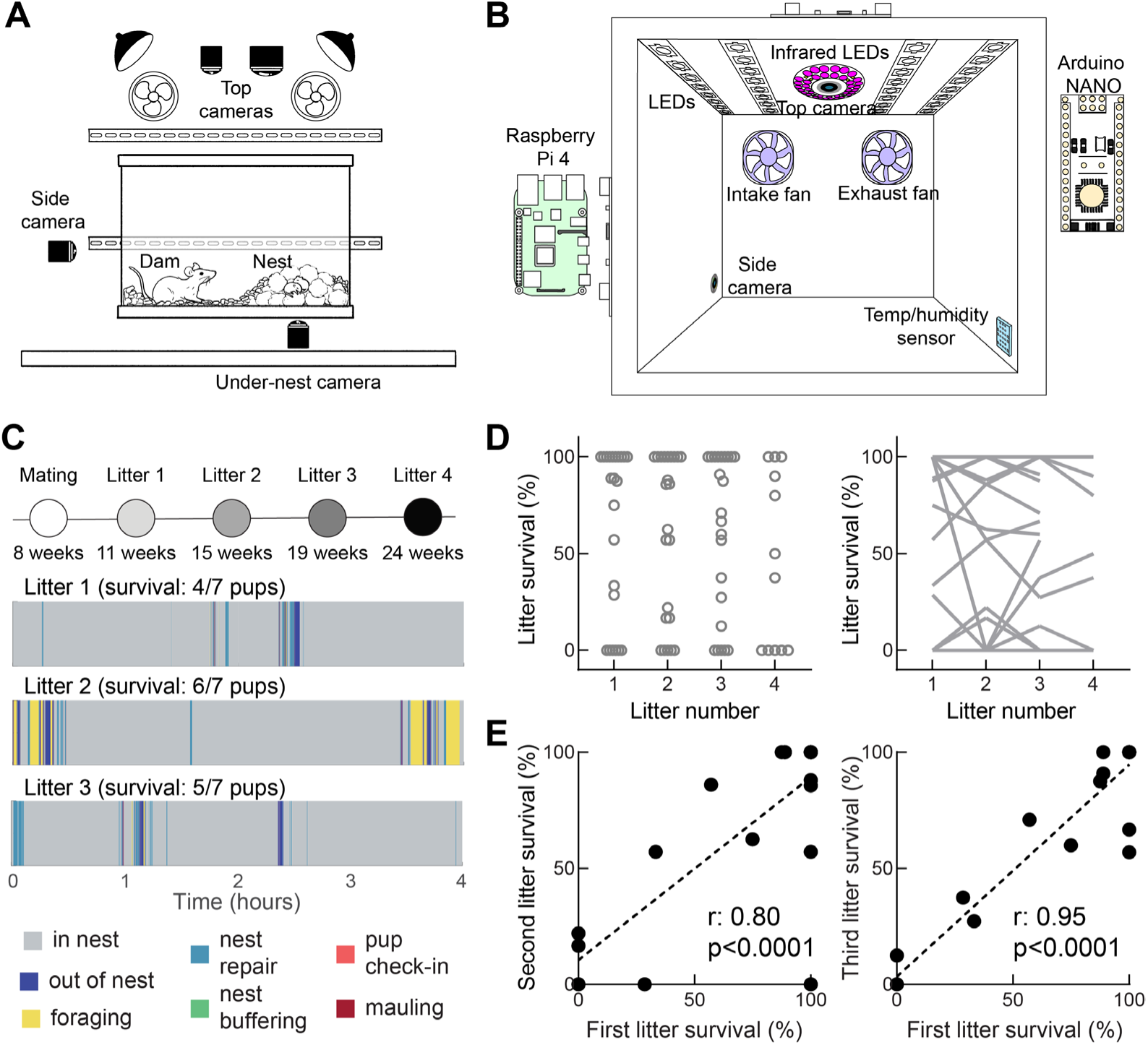
Continuous long-term recording of mouse maternal care across litters. (A) Setup for 24/7 continuous recording of homecage behaviors. A standard mouse homecage was placed within a custom enclosure containing cameras from top, side, and under the nest. (B) System for visible-light and thermal imaging with environmental controls. (C) Homecage behaviors were recorded over 3-4 consecutive litters, scored by automated methods (in/out of nest) and manually (foraging, nest repair, nest buffering with food pellets, pup and nest check-ins, mauling). Top, timeline of monitoring of individual animals (times are ages of the dams). Bottom, example ethograms showing several hours of spontaneous homecage behaviors for the same animal during litters 1, 2, and 3. (D) Litter survival at 24 hours post-parturition. Left, percent of pups surviving after the first postnatal day (N=22 dams). Each symbol represents fraction of litter survival for one dam over litters 1-4. 9/22 dams had 100% pup survival at litter 1, and 7/9 of those animals had 100% pup survival at litter 3. 6/22 dams had 0% pup survival at litter 1, and 5/6 of those animals had 0% pup survival at litter 3. Right, same data with lines connecting litter survival from each dam across litters. (E) Correlation of pup survival between litters. Left, litter 1 survival predicted litter 2 survival across dams (N=22, r=0.80, p<0.0001). Right, litter 1 survival predicted litter 3 survival (N=22, r=0.95, p<0.0001).

We monitored 65 single-housed wild-type C56Bl/6 female mice continuously 24 hours/day, 7 days/week over 3-4 months, from first mating and first litter through subsequent breeding and litters (**Fig. 1C**, top). We documented and scored many different maternal behaviors over the postnatal period, and focused on interactions with pups and the nest in the first four hours to days of each litter (**Fig. 1C**, bottom). This was due largely to surprisingly high pup mortality even in the protected conditions of the lab vivarium and standardized housing enclosures we developed.

We quantified pup survival from the moment of birth over the first 24 postnatal hours, in which most mortality was observed. Across the first cohort of 22 mice (**Fig. 1D**), the average pup survival rate from the first litter was 61.8±9.3%. However, pup survival was clearly bimodal: 9/22 first-time dams had 100% litter survival, and 6/22 dams had 0% litter survival (**Fig. 1D**, Litter 1). We refer to dams with ≥50% litter survival as ‘high-pup-survival’ dams, and dams with <50% pups surviving as ‘low-pup-survival’ dams.

We hypothesized that for low-pup-survival dams, litter survival rates might improve over subsequent litters as dams learned from experience. This was not the case. As shown in **Figure 1D**, pup mortality persisted across litters with 8/22 low-pup-survival dams for litter 2, 8/22 low-pup-survival dams for litter 3, and 6/11 low-pup-survival dams for litter 4 (**Fig. 1D**, left). With the exception of one dam, high pup survival persisted for high-pup-survival dams across litters (**Fig. 1D**, right). This meant that the pup survival rate of the first litter predicted with high accuracy pup survival in subsequent litters (**Fig. 1E**; N=22, r≥0.8, p<0.001).

### High-pup-survival dams spent more time investigating and repairing nests

Only a small fraction of pup mortality was due to aggression or mauling of pups (just 19/241 deceased pups from 5/65 dams; **Fig. 2A**, red). Instead, low-pup-survival dams neglected their pups and nests after initial nest-building was complete, and most pups died within the first four hours after birth (186/241 deceased pups; **Fig 2A**, black). Pup mortality seemed to result from hypothermia (**Supplementary Video 2**), based on thermal imaging of pup temperature (**Fig. 2B**).

**Figure 2.**
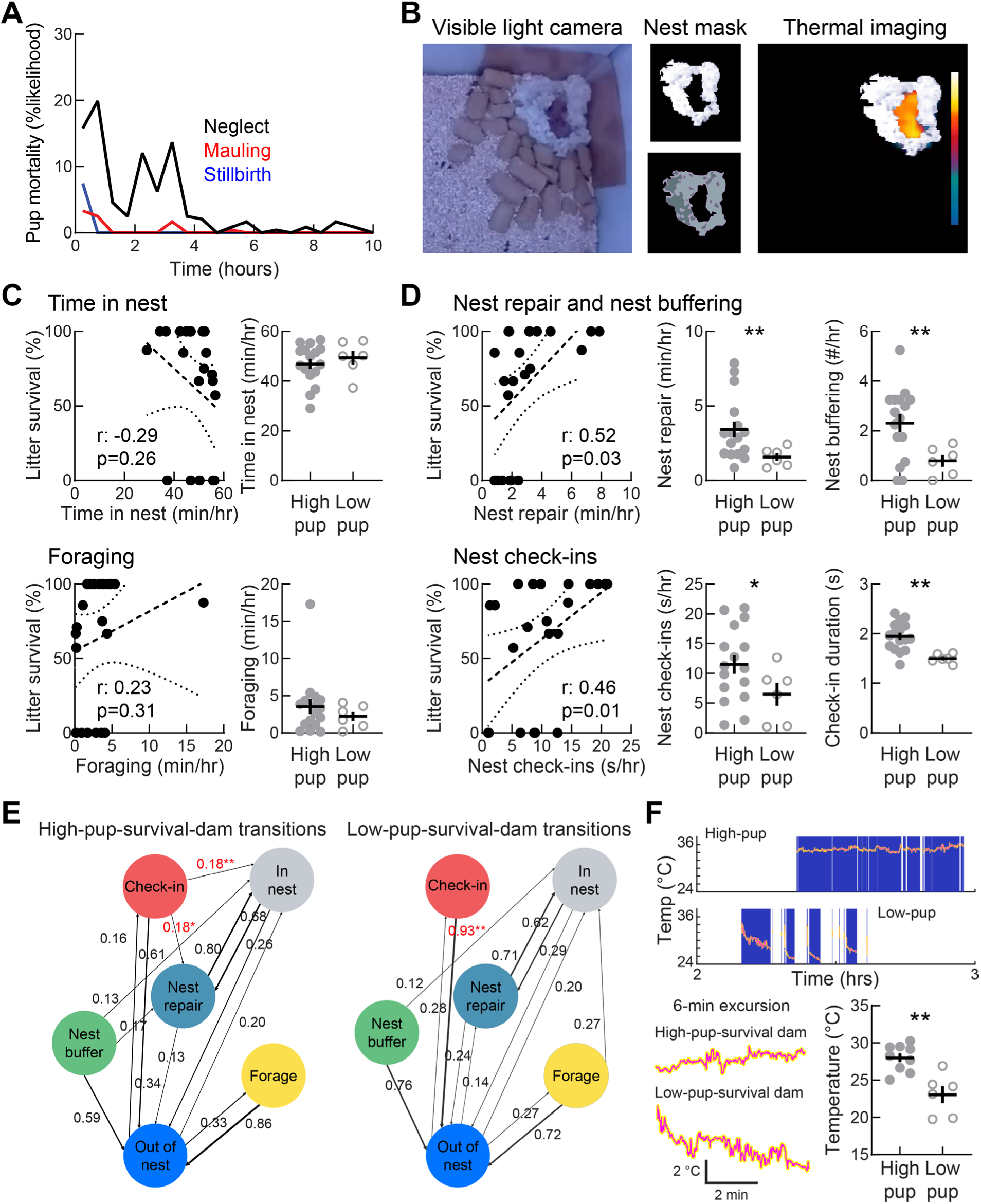
Nests must be rebuilt to ensure pup thermoregulation. (A) Causes of pup mortality in hours after birth. Most pups died from neglect (black, heat loss and/or lack of nutrients, 198/233 pups monitored over 10 hours) in the first four hours after birth (77.3% of mortality), rather than mauling (red, 19/233) or being stillborn (blue, 16/233 pups). (B) Procedure for monitoring pup temperature. Video frames were used to compute nest mask, registered with thermal imaging to isolate pup temperatures during periods when dam was off the nest. (C) Pup mortality was not correlated with time dam spent in nest nursing, grooming, or resting. Top left, no correlation across N=22 dams (r=-0.29, p=0.26). Top right, no difference of time in nest between high-pup-survival dams (46.9±2.0 min/hr) and low-pup-survival dams (49.4±2.8 min/hr, p=0.38, Benjamini-Hochberg multiple comparisons correction). Bottom left, no correlation with time spent foraging across N=22 dams (r=0.23, p=0.31). Bottom right no difference of time spent foraging between high-pup-survival dams (3.5±1.0 min/hr) and low-pup-survival dams (2.2±0.6 min/hr, p=0.13). (D) Pup survival was related to time spent repairing nest. Top left, correlation between time engaging in nest repair and pup survival (r=0.52, p=0.03). Top middle, high-pup-survival dams spent more time repairing nests (3.4±0.5 min/hr) compared to low-pup-survival dams (1.6±0.2 min/hr, p=0.005, Benjamini-Hochberg multiple comparisons correction). Top right, high-pup-survival dams buffered nest with food pellets (2.3±0.4 events/hr) more than low-pup-survival dams (0.8±0.2 events/hr, p=0.003). Bottom left, correlation between nest check-ins and pup survival (r=0.46, p=0.01). Bottom middle, high-pup-survival dams spent more time performing nest check-ins (11.5±1.6 sec/hr) compared to low-pup-survival dams (6.5±1.9 sec/hr, p=0.048). Bottom right, average check-in duration was longer for high-pup-survival dams (1.9±0.7 sec) compared to low-pup-survival dams (1.5±0.4 s/hr, p=0.00006). (E) Average frame-by-frame behavioral transition probabilities for high-and low-pup-survival dams, for time in nest, time out of nest, foraging, nest check-ins, nest buffering, and nest repair (not shown: self-loops and transitions occurring <10% of time). Transition probabilities after nest check-in were larger from check-in to nest repair for high-pup-survival dams (18.2±3.7%, N=16) than for low-pup-survival dams (5.2±3.0%, N=6, p=0.01, Student’s unpaired two-tailed t-test). Transition probabilities from check-in to entering nest were larger for high-pup-survival dams (18.1±4.1%) than for low-pup-survival dams (1.7±1.1%, p=0.001). Transition probabilities were lower from check-in to leaving nest for high-pup-survival dams (61.1±5.2%) than low-pup-survival dams (93.2±3.3%, p=0.00005). (F) Heat loss when low-pup-survival dams leave the nest. Top, pup temperature between hours 2-3 post-parturition for example high-and low-pup-survival dams. Blank periods are when dam was in nest (preventing measurement of pup temperature beneath her). High-pup-survival litter stayed warm due to nest insulation and exhibited minimal heat loss during dam excursions (temperature range 34-36 °C). Low-pup-survival litter rapidly cooled (dropping from 34 °C to 24-30 °C within minutes). Bottom left, zoom in on 6-minute excursion from traces above. Bottom right, summary of temperature drops across all litters with thermal imaging (high-pup-survival litters, N=9, lowest temperature: 26.0±1.8 °C; low-pup-survival litters, N=6, lowest temperature: 23.1±2.8 °C, p=0.005, Student’s unpaired two-tailed t-test).

We annotated videos in the first four hours post-partum using DeepLabCut to track dam and pup locations (**Fig. S1A**), as well as with manual annotation by trained observers blind to total pup mortality and if dams were assigned ‘high-’ or ‘low-pup-survival’ labels. Given the importance of nests to keep pups insulated, we focused on pup-and nest-directed behaviors: time of dam in nest (including nursing and grooming), time repairing nest, nest buffering, time spent foraging, time spent examining or checking-in on nest and pups, and the duration of each individual check-in event.

High-and low-pup-survival dams spent most time in the nest. However, time in the nest was not correlated with pup survival (**Fig. 2C**, top; ‘Time in nest’, N=22, r=-0.29, p=0.26; high-vs low-pup-survival p=0.38, Benjamini-Hochberg multiple comparisons correction). These behaviors included grooming, keeping pups warm, and nursing (**Fig. S1B**), but due to the challenges of accurately annotating dam and pup behaviors when pups were visually occluded on video, we combined these different behaviors into one category of ‘time in nest’. Similarly, time spent out of the nest foraging (eating, drinking, retrieving nest material) was also uncorrelated with pup mortality (**Fig. 2C**, bottom; ‘Foraging’, r=0.23, p=0.31; high-pup-vs low-pup-survival p=0.13).

Although high-pup-survival and low-pup-survival dams often initially built nests, a strong predictor of pup survival was amount of time the dam spent rebuilding the nest (**Fig. 2D**, top left and middle; ‘Nest repair’, r=0.52, p=0.03; high-vs low-pup-survival p=0.005). Low-pup-survival dams often initially spent some time nest-building, but then gave up and subsequently lost most or all of their offspring within hours. High-pup-survival dams continued to repair the nest especially after leaving and disrupting the nest integrity. Many high-pup-survival dams also stocked food pellets near the nest (**Fig. 2D**, top right; ‘Nest buffering’, high-vs low-pup-survival p=0.003), which bolstered nest wall integrity due to the mass of the food pellets.

Nest repair was related to high-pup-survival dams spending more time occasionally ‘checking-in’ on pups and nest, and high-pup-survival dams also spent more time for each check-in event (**Fig. 2D**, bottom; time spent on check-ins, r=0.46, p=0.01; high-pup-survival vs low-pup-survival p=0.048; average time per check-in, p=0.00006). Frame-by-frame quantification of behavioral patterns over the first four hours post-parturition revealed different transition probabilities from one form of behavior to another (e.g., in nest to out of nest, nest repair to nest buffer, etc.) for high-vs low-pup-survival dams (**Fig. 2E**, **Tables S1** and **S2**). While most of the behavioral transitions were similar and did not distinguish pup survival, the consequences of nest check-ins were significantly different. From each check-in, high-pup-survival dams tended to enter the nest or engage in nest repair immediately thereafter, whereas low-pup-survival dams remained outside the nest and neglected their pups (**Fig. 2E**, red transition probabilities).

To verify the importance of nest check-ins and nest repair for preventing hypothermia and promoting pup survival, we monitored pup temperature during periods when dams made excursions from the nest. This happened frequently as mother animals left to forage for food and water, and to regulate their own body temperature^3,29,30^. Nests from high-pup-survival dams kept pup temperature warmer and less variable over extended durations (**Fig. 2F**, top, bottom left). In contrast, pups from low-pup-survival dams rapidly cooled due to lack of a thermally-insulated nest, with temperatures dropping several degrees within minutes. Although they lived for a few hours, pups from low-pup-survival dams experienced more extreme temperature fluctuations and colder temperatures overall (**Fig. 2F**; high-pup-survival litters, N=9, lowest temperature: 26.0±1.8 °C; low-pup-survival litters, N=6, lowest temperature: 23.1±2.8 °C, p=0.005, Student’s unpaired two-tailed t-test). Thus the main feature distinguishing high-pup-survival from low-pup-survival dams was an increase in nest check-in time, leading to substantially more engagement with pups and nest thereafter to keep pups warm.

### Low-pup-survival dams can learn from co-housing with a high-pup-survival dam

We were surprised low-pup-survival dams did not increase attention to pups and time rebuilding nests across litters. Therefore, we next tested whether low-pup-survival dams would be unable to provide adequate caregiving under any conditions, or if they could become less neglectful if they had different parenting experiences. Previously we documented the emergence of alloparenting in initially-neglectful and pup-averse nulliparous (virgin) female C57Bl/6 mice, after being co-housed with experienced dams and litters^16,31^. Behavioral changes in the virgins were due to encouragement or coercion by the experienced dam, who shepherded or forced the virgin to spend time in the nest until eventually the virgin became a successful alloparent.

We hypothesized that similar parental behavior might emerge if we co-housed an initially low-pup-survival dam with an experienced dam and her litter. We identified a separate cohort of 43 low-pup-survival dams with 0% litter survival at litter 1 and ensured they were still low-pup-survival (<50% litter survival) for litter 2. Prior to mating these animals for litter 3, we co-housed some of them for 16-20 days with an experienced high-pup-survival dam and litter; some other animals were kept isolated for a similar duration. We then bred all the dams and monitored pup survival for litters 3 and 4 when they were returned to being singly-housed in our behavioral monitoring chambers (**Fig. 3A**), for a total of up to 32 weeks of continuous monitoring.

**Figure 3.**
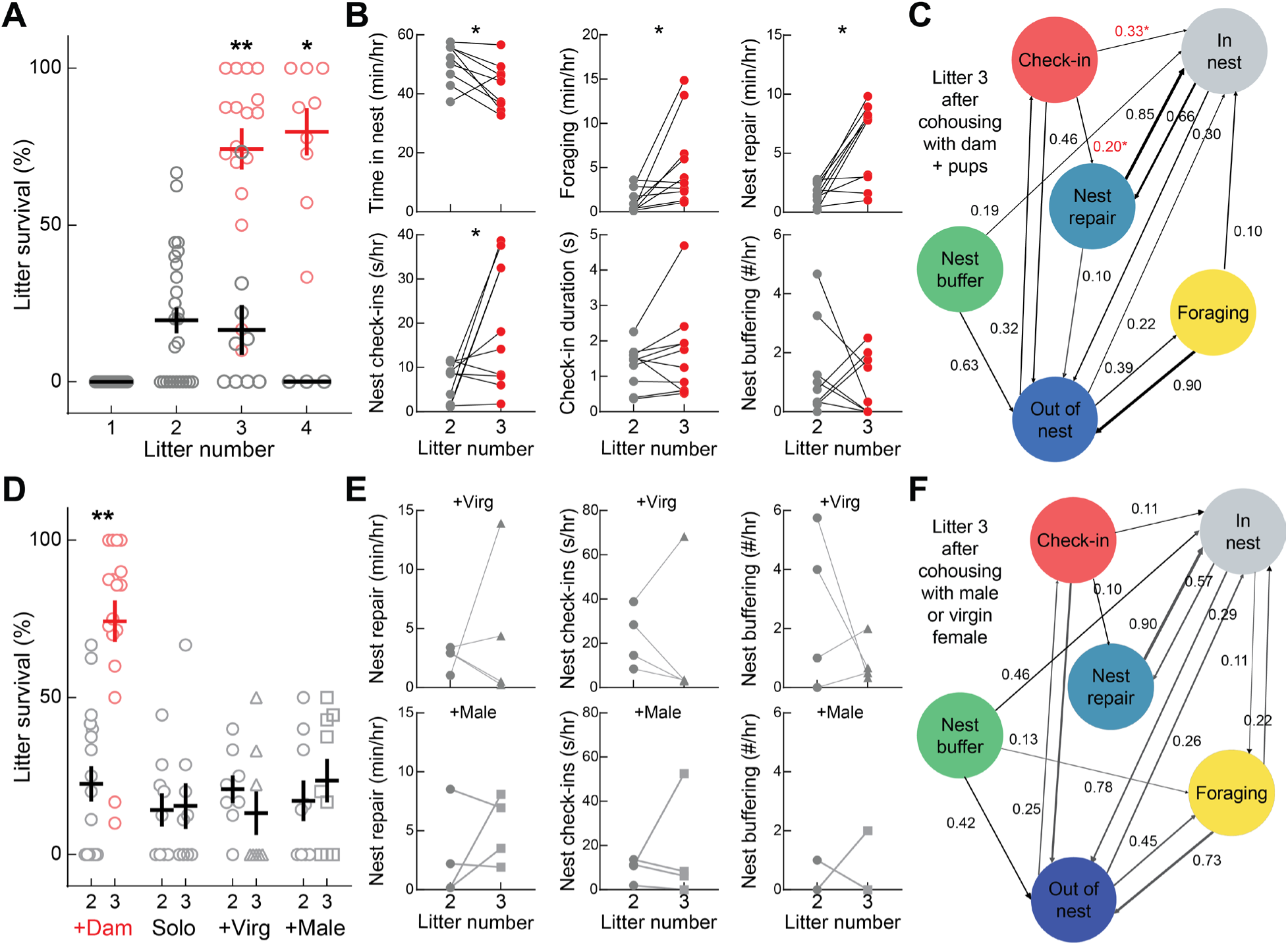
Co-housing with an experienced high-pup-survival dam improved maternal care. (A) Co-housing with an experienced high-pup-survival dam improved pup survival of initially low-pup-survival dams. Between litters 2 and 3, dams co-housed after litter 2 showed increased pup survival for litter 3 (74.2±6.6%, N=17) and litter 4 (79.7±7.5%, N=9). For the nine non-co-housed animals from the same cohort, litter survival remained low for litter 3 (15.4±7.3%, N=9, p<10^-5^ compared to litter 3 survival of co-housed animals, Student’s unpaired two-tailed t-test) and litter 4 (0%, N=3). Litter 2 survival was not different for the co-housed dams (22.5±5.7%) and the non-co-housed dams (14.2±5.3%; p=0.30, Student’s unpaired two-tailed t-test). (B) Some behaviors of initially-low-pup-survival dams were changed after temporary co-housing with an experienced high-pup-survival dam. Top left, time in nest decreased from litter 2 (50.4±2.2 min/hr, N=10) to litter 3 (42.7±2.5 min/hr, p=0.04, Student’s paired two-tailed t-test). Top middle, foraging time increased from litter 2 (1.3±0.4 min/hr) to litter 3 (5.5±1.5 min/hr, p=0.04). Top right, nest repair time increased from litter 2 (1.6±0.3 min/hr) to litter 3 (5.9±1.1 min/hr, p=0.02). Bottom left, time spent performing nest check-ins increased from litter 2 (6.6±1.3 sec/hr) to litter 3 (19.8±4.5 sec/hr, p=0.04). Bottom middle, check-in duration was unchanged from litter 2 (1.3±0.19 sec/event) to litter 3 (1.6±0.4 sec/event, p=0.29). Bottom right, nest buffering was unchanged from litter 2 (1.2±0.5 events/hr) to litter 3 (0.8±0.3 events/hr, p=0.54). (C) Average frame-by-frame behavioral transition probabilities for litter 3 of low-pup-survival dams co-housed with experienced dam and litter, for time in nest, time out of nest, foraging, nest check-ins, nest buffering, and nest repair (not shown: self-loops and transitions occurring <10% of time). Transition probabilities from nest check-in to nest repair increased between litter 2 (5.9±2.0%) compared to litter 3 (18.9±4.7%, p=0.039). Transition probabilities from nest check-in to nest entry increased from litter 2 (7.9±2.6%) to litter 3 (33.1±8.0%, p=0.023). (D) Initially-low-pup-survival dams co-housed with high-pup-survival dam and litter for 16-20 days between litters 2 and 3 led to higher pup survival at litter 3 (‘+Dam’, N=17, 74.2±6.6%) compared to litter 2 (22.5±5.7%, p<10^-4^, Student’s paired two-tailed t-test). There was no change in litter survival for dams not co-housed (‘Solo’, litter 2 survival: 14.2±5.3%, litter 3 survival: 15.4±7.3%, N=9, p=0.86), for dams co-housed with a nulliparous female (‘+Virg’, litter 2 survival: 20.8±4.4%, litter 3 survival: 13.2±7.0%, N=8, p=0.29), or for dams co-housed with the paternal male (‘+Male’, litter 2 survival: 17.1±6.5%, litter 3 survival: 23.5±6.9%, N=9, p=0.32). (E) Parental behaviors of low-pup-survival dams were not significantly changed from litter 2 to litter 3 after co-housing with a nulliparous female or the paternal male. (F) Average frame-by-frame behavioral transition probabilities for litter 3 of low-pup-survival dams co-housed with either a nulliparous female or the paternal male. None of the behavioral transitions were significantly different between litters 2 and 3 before vs after co-housing (for example: from check-in to nest repair, p=0.57; from check-in to nest entry, p=0.50).

Temporarily co-housing ‘low-pup-survival’ dams with another experienced mother led to much higher survival rates of litters 3 and 4, compared to dams which were isolated for 16-20 days (**Fig. 3A**; red, co-housed dams, litter 3 survival: 74.2±6.6%, N=17; black, isolate dams, litter 3 survival: 15.4±7.3%, N=9, p<10^-5^ compared to litter 3 survival of co-housed animals, Student’s unpaired two-tailed t-test). During the co-housing period, the low-pup-survival dams were initially pup-averse, but within hours to days they began interacting with pups and helped with childcare (**Supplementary Video 3**).

This was due to behaviors of the other experienced dam, who shepherded the low-pup-survival dam into the nest and demonstrated aspects of maternal care^16^. Shepherding led to more nest repair and nest check-ins for litter 3 as compared to litter 2 for these initially low-pup-survival dams (**Fig. 3B**; nest repair litter 3 vs litter 2, p=0.02; time spent on check-ins litter 3 vs litter 2, p=0.04, Student’s paired two-tailed t-test corrected for multiple comparisons with Benjamini-Hochberg). Total time in the nest decreased slightly and foraging time increased to a level similar to the high-pup-survival dams (**Fig. 3B**; time in nest litter 3 vs litter 2, p=0.04; foraging litter 3 vs litter 2, p=0.04). Not all behaviors became similar; low-pup-survival dams did not increase the duration of each nest check-in, nor did they increase nest buffering with food pellets or other environmental objects (**Fig. 3B**; nest check-in durations litter 3 vs litter 2, p=0.29; nest buffering litter 3 vs litter 2, p=0.54). This indicates that although sustained patterns of nest and pup monitoring are critical for infant survival, there are multiple different strategies for performing these tasks which each can lead to improved pup survival outcomes.

Frame-by-frame analyses of the behaviors of the low-pup-survival dams revealed that co-housed low-pup-survival dams also became similar to high-pup-survival animals for specific behavioral transitions. After nest check-ins, dams made more transitions to either nest repair or time in the nest (**Fig. 3C**; check-ins to nest repair litter 3 vs litter 2, p=0.04; check-ins to nest entry litter 3 vs litter 2, p=0.02; Student’s unpaired two-tailed t-test). These behaviors ensured that pups did not become hypothermic. Other behavioral patterns remained similar before and after co-housing (**Figs. 2E, 3C; Tables S3,4**).

Improvement of maternal care in low-pup-survival dams was not due to social buffering effects from co-housing with any other animal. Co-housing low-pup-survival dams between litters 2 and 3 with either a pup-naïve nulliparous (‘virgin’) female (N=8) or an experienced paternal male (N=9) did not improve pup survival for litter 3 (**Fig. 3D**; ‘+Virg’, co-housed with virgin female, litter 2 survival: 20.8±4.4%, litter 3 survival: 13.2±6.9%, N=8, p=0.29, paired two-tailed t-test; ‘+Male’, co-housed with paternal male, litter 2 survival: 17.1±6.5%, litter 3 survival: 23.5±6.9%, N=9, p=0.32). Co-housing in these cases occurred without pups, as none of the adult animals would be lactating. For 10 dams we were able to fully document the low-pup-survival dam behaviors before and after co-housing for litters 2 and 3. There was no difference in time spent engaged in parental behaviors before and after co-housing with either a virgin female (**Fig. 3E**, top, N=6) or paternal male (**Fig. 3E**, bottom, N=4).

Some of these dams that remained low-pup-survival spent substantial amounts of time checking in or repairing the nest, either before co-housing or after being paired with a companion male or nulliparous female. The total overall time spent engaging in different behaviors could sometimes appear similar to the behaviors of high-pup-survival dams. However, when we analyzed the frame-by-frame behavioral transitions during litter 3, we found no difference in the sequencing of when nest repair or nest entry occurred compared to litter 2 behavioral transitions (**Fig. 3F, Tables S5,6**). We conclude that total time with pups is insufficient. Instead, the critical factor is when these behaviors occur relative to pup needs, i.e., the contingencies or organization of parental caregiving behaviors critical for infant survival.

### Mouse ‘midwives’ aid delivery during dystocia and greatly increase mother-infant survival

Finally, we examined the broader utility of our system for phenotyping of maternal care and infant survival. As a test case, we studied OXTR-KO mice, given the interest in behaviors of animals lacking oxytocin receptors^21^. Specifically, we asked how animals lacking uterine contractions could have normal parturition^20^. Our video recordings from under pregnant dams during parturition revealed that some other animals actively engage in helping behaviors that mitigate the stresses of delivery and dystocia, and greatly improve maternal-infant survival (**Fig. 4A**).

**Figure 4.**
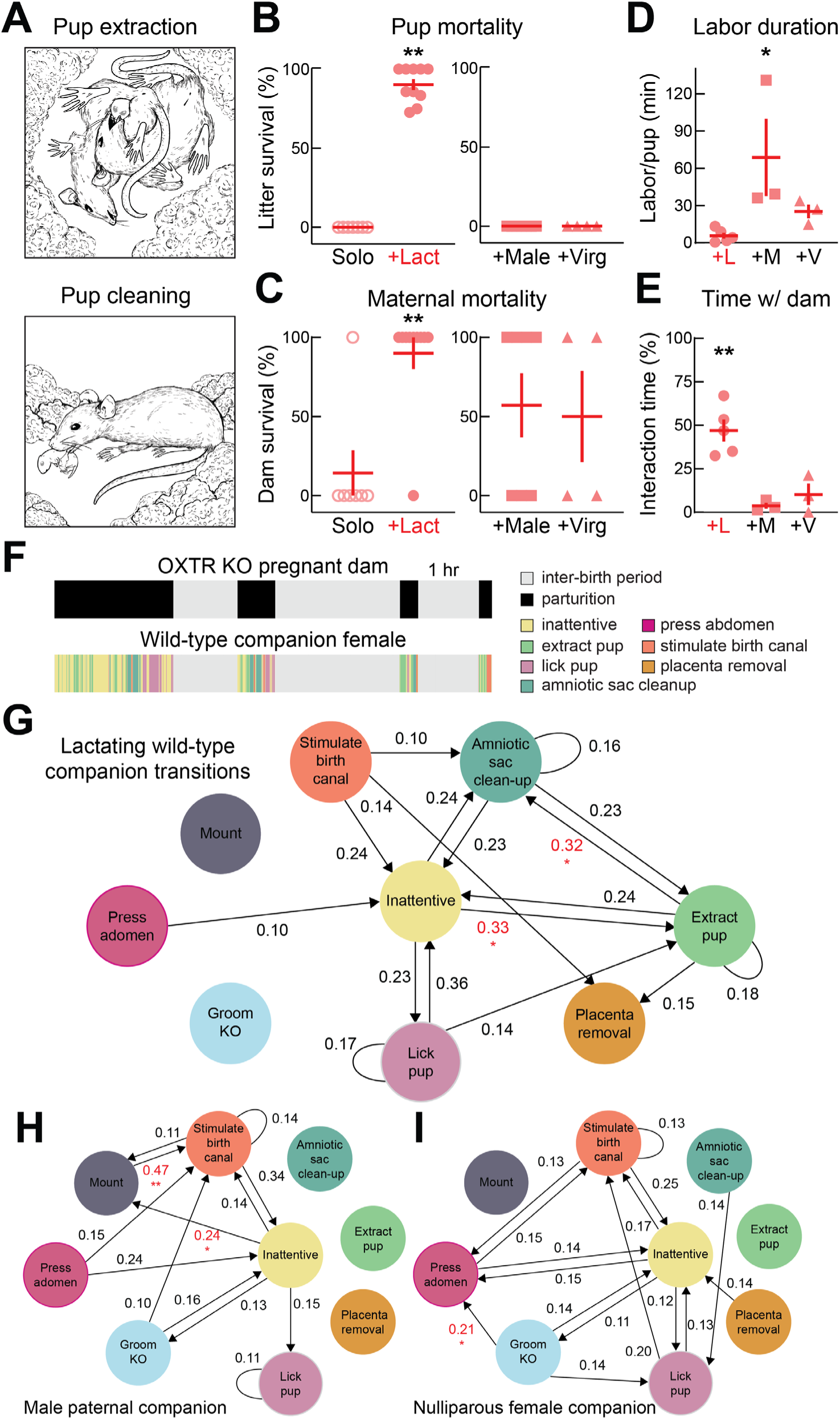
Co-housed companion animals improve OXTR-KO mother-infant survival. (A) Illustrations of two helping behaviors of lactating wild-type female companions during labor. Top, extracting stuck pups with mouth or paw. Bottom, cleaning pups of amniotic sac post-delivery. (B) Pup survival rates from primiparous OXTR-KO dams. Left, pregnant OXTR-KOs were singly housed (‘Solo’, N=7) or co-housed with a lactating wild-type female (‘+Lact’, N=10). Solo dams had 0% survival; co-housing with a lactating female led to 90.2±3.5% litter survival (p<10^-5^, Student’s unpaired two-tailed t-test). Right, pregnant OXTR-KOs co-housed with either the paternal wild-type male (‘+Male’, N=7) or a nulliparous wild-type female (‘+Virg’, N=4) had 0% pup survival. (C) Maternal survival of OXTR-KO dams. Left, pregnant OXTR-KO that were singly housed mostly died in labor (‘Solo’, 1/7 OXTR-KO dams survived labor). OXTR-KOs co-housed with lactating wild-type female almost all survived labor (‘+Lact’, 9/10 OXTR-KO dams survived, p<10^-5^, Fisher’s two-tailed exact test). Right, 4/7 pregnant OXTR-KOs co-housed with the paternal wild-type male survived; 2/4 pregnant OXTR-KOs co-housed with nulliparous wild-type female survived. (D) Labor duration was much shorter for pregnant OXTR-KO dams co-housed with a companion lactating female (average labor duration per pup: 5.8±2.4 min, N=5) or nulliparous female (average labor duration per pup: 25.3±5.5 min, N=3) compared to paternal male (average labor duration per pup: 68.8±31.1 min, N=3, p=0.034, ANOVA with Tukey’s multiple comparisons correction). (E) Total time attending to pregnant dam was much higher in companion lactating females (fraction of time spent interacting with pregnant OXTR-KO: 47.0±6.3%, N=5, p<0.0032, ANOVA with Tukey’s multiple comparisons correction) compared to paternal male (fraction of time spent interacting with pregnant OXTR-KO: 3.7±1.7%, N=3) or nulliparous female companions (fraction of time spent interacting with pregnant OXTR-KO: 10.3±6.2%, N=3). (F) Ethograms of behaviors of example co-housed pregnant OXTR-KO dam (black, time spent delivering each pup) and companion wild-type lactating female (pressing abdomen, stimulating birth canal, extracting pup with paw or mouth, licking pup, cleaning pup of amniotic sac, placenta removal). Note precise temporal synchronization of companion interactions only during moments of delivery for each pup. (G) Average frame-by-frame transition probabilities for behaviors of companion wild-type lactating females (N=5 adult pairs with 23 delivered pups). Periods of inattentiveness transitioned to pup extraction and pup extraction transitioned to pup cleaning significantly more for animals co-housed with lactating females than with male or nulliparous female companions. (H) Average frame-by-frame transition probabilities for behaviors of companion wild-type paternal males (N=3 adult pairs with 9 delivered pups). Periods of inattentiveness transitioned to mounting and mounting transitioned to birth canal stimulation significantly more than in female companions. (I) Average frame-by-frame transition probabilities for behaviors of companion wild-type nulliparous females (N=3 adult pairs with 7 delivered pups). Grooming the pregnant OXTR-KO dam transitioned to abdominal pressing significantly more than in lactating female or male companions.

The first cohort of OXTR-KO animals were singly-housed nulliparous KO females after the first round of mating and breeding. Pregnant animals turn in circles during labor, clearing the cage floor of bedding and facilitating unobstructed video recording from underneath the homecage during this time. To our surprise, singly-housed OXTR-KO animals labored for many hours or days, with pups becoming stuck in the birth canal and the pregnant animal attempting to pull them out with their mouth or paws, or mechanically push them out by abdominal pressure (**Supplementary Video 4**). The prolonged and difficult labor (‘dystocia’) prevented them from cleaning pups of the amniotic sac to stimulate breathing^32^. As a consequence of hypoxia and lack of milk ejection in OXTR-KO females^20^, all pups born to singly-housed OXTR-KO females died (**Fig. 4B**, left; ‘Solo’, N=7, 0% pup survival for all litters). Nearly all the singly-housed pregnant OXTR-KO females died as well (**Fig. 4C**, left; ‘Solo’, 6/7 OXTR-KO females expired).

Much mouse breeding occurs in group-housed cages (e.g., for harem breeding and to reduce the number of animal cages and cage costs). In addition, because OXTR-KO mice do not have milk ejection during nursing^20^, pups must be co-housed with a lactating female to survive. Thus we tested OXTR-KO females co-housed with a lactating wild-type female companion (‘Lact’), close to term or recently delivered, i.e., dams at gestation day 19 (GD19) to P0. The pregnant OXTR-KO female had prolonged and difficult labor, but when the first pup crowned and got stuck in the birth canal, we observed remarkable changes in behavior of the companion lactating female mouse. The other female approached the OXTR-KO mouse in dystocia, and acted as a ‘midwife’, helping to remove stuck pups with her mouth and paws, and cleaning pups of the amniotic sac so they could breathe (**Supplementary Video 5**). These helping behaviors seemed vital to maternal-infant survival. In all cases, pup mortality was drastically reduced and litter survival rates were quite high (**Fig. 4B**, left; ‘+Lact’, 90.2±3.5% pup survival per litter, N=10 litters, p<10^-5^ compared to singly-housed OXTR-KO litters, Student’s two-tailed unpaired t-test). 9/10 co-housed pregnant OXTR-KO females also survived labor, compared to 1/7 singly-housed OXTR-KOs (**Fig. 4C**, left, ‘+Lact’, p<0.003, Fisher’s two-tailed exact test).

We asked if other adult animals co-housed with the pregnant OXTR-KO interacted with the female in labor. If the companion animal was either the paternal male (‘Male’) or a nulliparous wild-type female (‘Virg’), all pups died (**Fig. 4B**, right), although maternal survival somewhat improved (**Fig. 4C**, right; ‘+Male’, 4/7 pregnant OXTR-KOs survived; +Virg’, 2/4 pregnant OXTR-KOs survived). It seemed as though each type of companion was engaging with the pregnant OXTR-KO dam in different ways, as the total amount of time spent delivering each pup differed, with short delivery times for OXTR-KO dams paired with lactating wild-type females (**Fig. 4D**; ‘+L’, delivery time per pup: 5.8±2.4 min, N=5), and much longer average times per pup for dams paired with paternal males (**Fig. 4D**; ‘+M’, delivery time per pup: 68.8±31.1 min, N=3, p=0.034 compared to labor time with lactating companion, ANOVA with Tukey’s correction for multiple comparisons). We also noticed a substantial difference in the total time the lactating wild-type female companion was interacting with the OXTR-KO during parturition (**Fig. 4E**; wild-type lactating companion, ‘+L’, interacting with pregnant dam 47.0±6.3% of total time over parturition, N=5, p<0.003 compared to male or nulliparous female companion interaction times; wild-type male companion, ‘+M’, interaction time 3.7±1.7% of total time; wild-type nulliparous female companion, ‘+V’, 10.3±6.2% of total time).

To determine the differences of interactions between lactating female companions and males or nulliparous females with the pregnant dam, we performed frame-by-frame analysis of companion animal behavior throughout parturition, with synchronized video feeds from the top, side, and underneath the pregnant dam. Analysis was performed for 23 pups successfully born to five OXTR-KO females co-housed with a lactating wild-type female, nine pups born to three OXTR-KO females co-housed with a paternal male, and seven pups born to three OXTR-KO females co-housed with a nulliparous female companion. An example ethogram for an OXTR-KO pregnant dam paired with a wild-type lactating female is shown in **Figure 4F** for one hour. During each pup birthing event, the lactating female companion animal engaged in several different behaviors, marked by brief inattentive periods before resuming interaction with the pregnant dam. These interactions only occurred during birthing, and the lactating female otherwise did not interact with the pregnant dam when she was not in active labor with the next pup. The relative times spent performing each behavior during birthing is quantified in **Figure S2**.

We examined the temporal organization of different behaviors enacted by wild-type lactating females (**Fig. 4G, Table S7**, N=5 dams, 23 pups), paternal males (**Fig. 4H, Table S8**, N=3 dams, 9 pups), or nulliparous females (**Fig. 4I, Table S9**, N=3 dams, 7 pups) towards the pregnant dam. Although there was considerable variability in the patterns of interactions for each delivered pup and across animals, a few general principles emerged that related to the degree of mother-infant survival. Lactating female companions frequently stimulated the birth canal, attempted to extract stuck pups with their mouth or paw, cleaned pups of the amniotic sac, and removed and scavenged the placenta after each birthing (**Fig. S2**). Lactating females transitioned from periods of inactivity to pup extraction (32.9±7.6% transition probability, p<0.03 compared to paternal and nulliparous companions), and transitioned from pup extraction to amniotic sac cleaning (31.7±7.5% transition probability, p<0.04) much more frequently, as these events were never observed in males or nulliparous females (**Fig. 4G-I**). This attention and help towards the mothers likely improved maternal survival of these co-housed animals, and subsequent attention towards each pup once removed seemed to lead to the much higher (non-zero) pup survival rates.

In contrast, males and nulliparous females engaged in other behaviors, which helped keep some of the OXTR-KO dams alive, but still led to 100% pup mortality. From inattentive periods, males mounted the pregnant dam (**Fig. 4H**; 23.9±7.2%, p<0.02) then stimulated the birth canal (**Fig. 4H**; 46.6±9.7%, p<0.0005), and both behaviors could help push out pups. Nulliparous females did not mount the dams, but groomed them followed by abdominal pressure which helped extract pups (**Fig. 4I**; 21.4±5.6%, p<0.02). However, once a pup was birthed, it was neglected and subsequently perished. Thus other adults can aid in birthing, but the precise details and temporal sequencing of when and what the companion does has major impact on outcomes for mother and offspring.

## Discussion

Continuous video monitoring revealed critical behaviors and organized behavioral sequences that differentiated high-from low-pup-survival dams. Some of the most important differences were not due to the total duration or frequency of caregiving behaviors, but rather to the temporal organization and embedding within the larger ongoing context. Nest check-ins were found to be critical divergence points between the two groups: while low-pup-survival dams tended to remain outside after nest check-ins, high-pup-survival dams instead transitioned to move into the nest or engage in nest repair. This resulted in shorter but more frequent nest visits, avoiding prolonged absences and maintaining regular pup contact. These observations show how the structural and temporal variation of maternal behavior in early postpartum period shapes pup outcomes, specifically by helping to keep pups warm especially when caretakers are out of the nesr.

Our approach highlights the natural variability of maternal behavior, enabling a more direct comparison with the human parental experience. A considerable fraction of dams were low-pup-survival or even 0% survival, losing every infant across consecutive litters. This indicates that maternal neglect in mice is not an anomalous occurrence in full distributions of parental strategies and outcomes for infant health and mortality, but was recurrent and predictable. Differences in behaviors between the low-pup-survival dams and the high-pup-survival dams were unlikely to be due to external stressors, as our behavioral boxes were designed to normalize environmental factors across animals. While some past studies of parental behavior may have excluded low-pup-survival animals for various reasons, we believe it will be important in the future to examine the neural differences between low-pup-survival and high-pup-survival dams, which might aid translational efforts to diagnose, treat, and understand peripartum disorders.

Low-pup-survival and high-pup-survival maternal profiles might have some degree of genetic basis, but these phenotypes are not static or fixed traits-they can be easily, rapidly, and dramatically transformed by social experience. This behavioral plasticity for parenting behavior can be seen in those low-pup-survival dams that became successful caregivers in subsequent litters after being cohoused with an experienced dam and her pups. The success of intervention and prevention programs that teach human parents to detect and respond to social signals from children underscores the primary importance of teaching, learning, and training for promoting successful parenting and reducing child neglect or maltreatment^33–35^.

We discovered that pregnant and post-parturient wild-type females spontaneously assisted OXTR-KO mice in active labor, reducing the risk of dystocia and greatly enhancing pup and maternal survival outcomes. These findings are consistent with previous work in humans and other primates where birth assistance has been associated with reduced neonatal and maternal mortality^36^. This raises the possibility that the neuroendocrine adaptations associated with gestation may contribute to the increase of responsiveness to parturition-related signals in conspecifics, shaping and facilitating the assistive midwifery response. Even though males and virgin females did not engage in behaviors to directly facilitate the birthing process, they did interact with the parturient OXTR-KO and appeared to respond to its discomfort. Males exhibited an increase in mounting behavior, which may have indirectly facilitated labor by exerting pressure on the abdomen aiding pup delivery. Other interactions such as allolicking or allogrooming may represent forms of consolation or helping behavior, as previously documented in rodents responding to a conspecific in distress^37–44^.

Our findings indicate that even under the controlled and protected conditions of a laboratory vivarium, infant care can be immensely challenging and taxing, requiring sustained effort on the part of caregivers to ensure pups are fed and warm. Under the conditions of our environmentally-controlled home cages, approximately a third to a half of C57Bl/6 mouse mothers were observed to lose some to all of their litters due to neglect of pups and nest. The nest structure was quite dynamic, needing to be repeatedly adjusted or rebuilt due to the movement of the mother in and out of the nest, and the motion of the pups. Thermal imaging was important to relate this aspect of parental behavior to litter survival, as pups rapidly dropped in temperature without a well-constructed insulating nest and died within minutes. Given the importance of nest repair for pup survival, it will be interesting to determine how parental animals sense pup need and nest status, potentially leading to interruption of ongoing behaviors to seek out nesting material or attempt to fix the nest. Parental animals might be able to rely solely on active pup cues such as distress calls to gauge pup and nest state; alternatively, experienced parents might have strategies or an internal model for ensuring nest integrity.

One potential concern is that the extent of pup mortality and relatively high prevalence of low-pup-survival dams is a consequence of sustained single-adult housing, especially combined with the stress of childcare. A related caveat is that our studies were performed exclusively in mice from a single genetic background. Even though C57BL/6 mice provide a controlled baseline, it is believed that maternal nurturance and other related behaviors vary across strains^45^. Bendesky et al. (ref. 46) identified differences in vasopressin gene expression as leading to variation in nest-building behavior in species of *Peromyscus* mice. Vasopressin is a nine-amino-acid neuropeptide hormone related in structure and function to oxytocin^14^, and it is interesting to speculate on why pituitary peptides important or required for physiological function and reproduction would also be important for social cognition and parental behavior, and thus the survival of the species. Our results discovered here suggest that the demands of single parenting are so intense and stressful, that unless animals are able to obtain help and support for labor assistance and child rearing, it is probable that some to all offspring will quickly perish.

Human maternal mortality doubled in the United States across demographics between 1999-2019 (ref. 47). Human infant mortality historically has been quite high; it is estimated that until the 1950s, about 1 in 4 infants did not survive to their first year, and roughly half of children did not survive to age 15 (ref. 48). With advances in science, medicine, and cultural mechanisms for cooperative parenting, infant mortality is now much lower in many countries, although remains non-zero everywhere. For mice, most wild species are thought to live in multi-female colonies^8^ where perhaps many different adults can participate in childcare, tending to pups and nests, and shepherd or help other inexperienced adults take care of offspring. To date little is known about the fine-scale adult social dynamics of colony organization and how this relates to communal child-care. The system we have developed enables a systematic and parametric exploration of different variables related to social structure and housing conditions, and promises to reveal much more about the family life of lab mice and other species.

## Resource availability

### Lead contact

Requests for further information and resources should be directed to and will be fulfilled by the lead contact, Robert C. Froemke.

### Materials availability

This study did not generate new unique reagents.

### Data and code availability

Data that support the findings of this study are available on Databrary (www.databrary.org). Custom code are available on Github, publicly available as of the date of publication. Any additional information required to reanalyze the data reported in this paper is available from the lead contact upon request.

## Acknowledgements

We thank D. Lin and R.M. Sullivan for comments, discussions, and technical assistance. This work was funded by the BRAIN Initiative (NS107616 to R.C.F. and A.M.), NICHD (HD088411 to R.C.F., HD094830 to K.E.A.), NIMH (MH019524 to J.L.M.), NINDS (NS138066 to R.C.F., NS139546 to A.C.), Pew Charitable Trusts (to R.C.F.), and NSF (graduate research award to L.S.). Artwork in **Figures 1A** and **4A** was by Shari E. Ross.

## Author contributions

L.S., A.M., and R.C.F. designed and built the 24/7 monitoring system. L.S. and V.J.I. performed behavioral experiments. L.S., V.J.I., D.M.G.S., R.H., A.C., J.L.M., G.K., S.S., D.A., M.K., S.B.W., P.L., and R.C.F. performed behavioral analysis. J.L.M., and L.K. provided technical assistance with animal breeding and housing. K.E.A. provided analysis tools and data archiving. L.S. and R.C.F. designed the study and wrote the paper; L.S., R.C.F., K.E.A., and A.M. discussed and edited the paper.

## Declaration of interests

The authors declare no competing interests.

## Materials and Methods

All procedures were approved under NYU Grossman School of Medicine IACUC protocols.

## System for Long-Term 24/7 Behavioral Analysis

For long-term monitoring of the acquisition and expression of maternal behavior in single-housed animals, we used 8-10 week-old C57BL/6 naïve virgin female mice. No statistical methods were used to predetermine sample size. The experiments were not randomized. The behavioral raters were blinded to litter survival rate, allocation during co-housing, and outcome assessment.

A nulliparous female mouse was housed alone in a 27.94 × 27.94 × 27.94 cm plexiglass homecage covered with plentiful bedding material, food pellets, hydrogel and/or a water bottle (refreshed every 3-5 days), and one Nestlet. Homecages were contained within custom-designed environmental control boxes (35 x 35 x 35 cm) equipped with 2-5 camera modules, each outfitted with 160° adjustable night vision wide-angle lens and individually connected to Raspberry Pi (RPi) circuits to record from the top, side, and underneath the nest area. A separate thermal imaging camera was positioned above the nest and connected to a Windows PC. Conditions of each enclosure were automatically monitored and controlled by USB-powered Arduino Uno or Nano microcontrollers connected to sensors and actuators that maintained a consistent internal environment. Energy efficient, low voltage, individually programmable LED light-strips and four photosensitive LED infrared light rings were installed to provide a naturalistic light/dark cycle (12h-12h). A DHT11 sensor sampled the temperature and humidity of the box at 1-second intervals, and based on these readings the microcontroller adjusted the speeds of two silent case fans (intake/exhaust) to maintain the enclosure temperature at ∼26-28°C.

We began by recording continuous video of the nulliparous female while housed alone for up to 5 days, before introducing an experienced C57BL/6 male mouse for breeding afterwards. The male mouse was removed from the enclosure typically after 24-48 hours, and successful intercourse was confirmed through visual examination of video footage and the presence of a vaginal plug. We closely monitored the female mouse for physical and behavioral changes typical of a gravid animal. If by day 10 post-mating, the female mouse did not show weight-gain or weight-redistribution, a new experienced male was introduced in the homecage for mating. Dams were allowed to stay with the pups until weaning on P21 in the same homecage. After that, the female was placed back in the enclosure for a few days and mated again, following the same procedure as with the first litter. This was performed for four consecutive litters.

### Video Acquisition

Video recordings were acquired from 1-3 cameras (OV5647 sensor) sensitive to infrared light and outfitted with 160 FOV wide-angle lenses and placed either above the enclosure (overhead view), perpendicular to the nest (nest view), or below the nest (bottom-up view). Each camera was connected to a Raspberry Pi 4 single board computer^49^ that captured, streamed, and saved uninterrupted video from the enclosures (30 frames per second) to a local server using the Python library OpenCV. For some animals, videography was performed continuously at all times up to weaning of the third litter; for other animals, we focused video recording on 3-5 days before birth up until P5.

### Behavioral Analysis

For analysis of video data, data streams were first aligned using a custom MATLAB algorithm to synchronize the RPi video and thermal recordings by matching respective timestamp data. Both devices were connected to the same network, enabling information from the CSV files to be used for temporal alignment with a high degree of accuracy between the two data streams. Due to the differing frame rates between the RPi (30 frames/sec) and thermal camera recordings (5-15 frames/sec), temperature data were interpolated to match the higher frame rate of the RPi videos. This process of interpolation ensured that each video frame was assigned a corresponding temperature reading, enabling consistent synchronization of the visual and thermal data streams for analysis.

For pose estimation and animal tracking, top-down video recordings were processed using either an NVIDIA GeForce 3090 RTX or an NVIDIA Quadro RTX 5000 GPU running DeepLabCut^50,51^. We used markerless body part tracking of mice and pups in their homecages using three different networks depending on conditions (tracking single dams, co-housed dyads, or pups). Across all conditions, Resnet-50-based and dlcrnet_ms5-based neural networks were trained to identify key body parts for tracking, and the output saved in CSV format. For single-housed dams, the network was trained using 3162 manually-labeled frames (2238 manually-labeled frames with mice, 924 frames of empty cage) from 45 adult C57BL/6J dams. Labeled body parts included nose, left ear, right ear, neck, left shoulder, right shoulder, body center, left hip, right hip, tail base, and tail tip. The output of this neural network was processed with a custom MATLAB script to calculate the time spent in the nest or position relative to the nest, and to map behavioral trajectories.

A second DeepLabCut network was trained to track two adult co-housed mice, using the same set of body part labels as the single animal condition. 531 manually-labeled frames from 28 adult mice were used to train and refine the network. Output files were processed through a MATLAB script to analyze the position of each mouse relative to a region of interest (ROI) and calculate the distance between the two mice over time.

A third network was trained to track the body parts of pups from P0-P10. To account for developmental changes occurring over these ten days (e.g., size, color, proportions) 240 images from pups within this range were labeled. The labeled pup body parts differed from the ones in the adult networks, as these markers were better suited for the training of the network because of their inter-body part distance and morphology. These segments included the nose, left paw, right paw, left foot, right foot, and tail tip. Videos from both the single-housed and co-housed conditions were tracked, and outputs were processed with a custom MATLAB script that identified when the pups were out of the nest.

For behavioral coding and ethogram generation of singly-housed dams, videos over the first 4 hours from the birth of the last pup in the litter were manually annotated using Datavyu^52^. The timepoint at which the last pup was born was determined by visually inspecting the three camera views: top-down, side, and under the nest. After confirming the last birth, top-down videos were used as the primary source for behavioral coding, with side and under-nest perspectives only used when behavior could not be easily identified from the top-down view due to occlusions. We did not include behavioral transitions occurring <10% of the time or self-loops, as we separately computed and analyzed the total times spent engaged in each behavior.

Ethograms included the following behaviors: dam in the nest, dam out of the nest, foraging, nest building and repair, pup check-ins, and pup mauling. Dam in nest or out of nest refer to the position of the dam rather than a specific behavior, as some actions such as nursing or pup interactions were often occluded from view of any of the cameras. Specifically, these behaviors were defined as follows: Dam in nest: The mother was physically present in the nest, in contact with or in close proximity to her pups. This state encompasses various pup-directed behaviors such as nursing, licking/grooming, and pup retrieval. The location of the nest was extracted with custom Matlab code using image segmentation. Nest entry time was defined as the moment when the two front paws entered the nest cavity.

Dam out of nest: The mother was physically absent from the nest and not in contact with her pups. This state includes time spent exploring the cage, self-grooming, and resting away from the nest.

Foraging: The mother was actively searching for, manipulating, or consuming food or water. Foraging bouts were typically characterized by directed locomotion towards food sources, sniffing, and handling of food pellets or the water bottle spout.

Nest building and repair: The dam was actively manipulating and rearranging the already shredded cotton nest structure. This behavior is characterized by repeated shredding with the mouth, pawing, and displacing the nesting material.

Pup check-ins: These were brief, rapid returns to the nest by the mother during periods of nest absence, often accompanied by sniffing, orienting, nudging, or brief contact with pups before leaving again. These bouts were typically quite short (lasting a few seconds in duration) and did not involve prolonged nest occupancy or pup interaction.

Nest buffering: The mother pushed food pellets towards the perimeter of the nest. This behavior was distinct from typical hoarding in that the food was not consumed or stored for later use, but rather appeared to serve a structural or insulating function.

Pup mauling: Active killing of the pup. Dam manipulates a moving pup with her paws and mouth and chews on it. Typically, blood can be observed during these episodes and the pup stops moving shortly after. These events were extremely rare.

Between 2-6 independent coders identified each of these behaviors frame by frame and scored the videos blind to litter number and survival rate. The results from each rater were compared, compiled, and cross-validated. Cohen’s kappa^53^ was used to calculate inter-rater similarity across different observers scoring behaviors from the same video segments, and ranged from 0.75 to 0.97.

### Thermal Imaging

For thermal imaging, we recorded continuous thermograms with a thermal camera (FLIR C3) placed above the nest, streaming temperature data (5-15 frames/sec) to a Windows PC through a USB cable. Thermographic data were captured with the FLIR Tools software, timestamped, and saved as a csv file.

We synchronized the thermograms and the video recordings post-hoc using timestamp information. We isolated the episodes in which the dam was out of the nest, identified the video frame and thermogram corresponding to the onset of the episode, located the nest opening using image segmentation, and then extracted and averaged the corresponding pixel temperature information from that ROI. The selected frames were processed using custom MATLAB scripts. Each frame was first converted to grayscale and denoised using a convolutional neural network. Contrast was enhanced using histogram equalization. The image was then converted to HSV color space to improve the distinction between the nest and background. Texture analysis was applied to emphasize structural details in the nest, and a flood-fill algorithm was employed to accurately isolate the nest from surrounding areas. We saved these frame-by-frame average traces as csv files. For pup temperature analysis, we identified episodes in which pups were out of the nest and followed a similar procedure to the nest temperature analysis focusing on the out-of-nest pup as our ROI.

### Litter Mortality

We monitored both the videos and the thermograms daily to establish how many pups were born alive, how many were stillbirths, and how many were cannibalized. We used color, temperature, movement, and physical integrity to determine if pups were alive. We calculated the litter survival rate as surviving pups on P5 divided by the total number of pups born alive.

### Co-Housing

Dams with 0% litter survival during their first litter and less than 50% survival during their second litter were co-housed either with an experienced high-pup-survival dam with her pups (∼P5 and older), a virgin, a paternal male, or left in isolation. Co-housing was performed for 16-20 days, before being mated a third and fourth time.

### Parturition and Assisted Labor

Individual OXTR-KO females were placed in a new breeding cage with an experienced wild-type male for mating. Females were checked daily for copulation plugs and assigned GD 0 (gestation day 0) if the plug was present. Males remained in the cage until a plug was seen but were taken out no later than seven days, whichever occurred first. On day 15, OXTR-KO females were visually inspected, and successful pregnancy was determined by increase in abdominal size^54^. The OXTR-KO was included in the study only if it was pregnant. On day 18, the OXTR-KOs were moved to the behavioral enclosure and singly-housed for observation and recording.

For studies involving co-housing with a companion animal during parturition, individual wild type females were placed in a new breeding cage in the vivarium with an age-matched female OXTR-KO and an experienced wild-type male for mating as above. Females were checked daily for copulation plugs and assigned GD 0 if the plug was present. The pairs of OXTR-KO and wild-type female were included in the study if both animals were pregnant. On day 18, both animals were moved to the behavioral enclosure for observation and recording. The tails of each of the pregnant mice were labeled with different colors and patterns to facilitate identification. Wild-type females were at similar gestational stages as OXTR-KOs; one animal gave birth before the companion OXTR KO female, and two wild-type companion females a few hours later the same day.

In some experiments, the wild-type companion was a nulliparous female. Immediately after mating the OXTR-KO, the mated female was placed in a new cage with an age-matched wild-type nulliparous female. In other experiments, the wild-type companion was the male mating partner, who remained in the cage after mating and throughout gestation. On GD 18, both animals were moved to the behavioral enclosure for observation and recording.

Starting on GD 18 when animals were moved to the behavioral enclosure, we inspected the video feed every 30 minutes for signs of labor (e.g., hunching, restlessness), and re-positioned the under-nest camera as needed to the location at start of parturition. Videos containing births of the first 4 pups were manually annotated using Datavyu. We phenotyped the behaviors of the cohoused mouse during active labor, from birth initiation to placental expulsion/removal. Birth initiation was determined visually by inspecting the dilation of the birth canal of the parturient OXTR-KO female, and labor concluded when the placenta was self-expelled or removed by the cohoused mouse. We classified the behaviors into attentive (birth assistance behaviors, social interactions with the mouse in labor) or inattentive.

Birth assistance behaviors include:

Amniotic sac cleanup: The companion mouse removes the amniotic membrane from a partially or fully delivered pup by using its mouth to tear, peel, or handle the sac, primarily targeting the pup head first and then the body. Licking behaviors may be observed during sac removal, contributing to the clearing of fluid from the pup’s airways.

Extract pup: The companion mouse aids in the removal of a partially delivered pup from the birth canal by grasping the pup with the forepaws or mouth and exerting outward pulling force. The companion mouse may alternate between paw-and mouth-based extraction methods and adjust grip location along the pup’s body.

Lick pup: The companion mouse licks the body of a partially or fully delivered pup. This behavior requires the amniotic sac involves repeated tongue movements directed at the head, body, and limbs of the pup. Licking is sustained for variable durations and may involve specific attention to the pup ano-genital region.

Placenta removal: The companion mouse interacts with the placenta of the pregnant female by gripping it with its forepaws or mouth and applying force to extract it from the birth canal after a pup has been delivered. Following removal, the midwife may ingest the placenta in its entirety or in fragments^55^.

Press abdomen: The companion mouse applies mechanical pressure to the abdominal region of the dam in labor. Force can be applied in different ways, including using forepaws to press directly on the abdomen, positioning head or snout under dam to exert localized pressure, or applying full body weight by climbing onto the back of the pregnant animal and positioning itself dorsally, which causes the compression of the pregnant animal’s abdomen against the cage floor, resulting in additional mechanical force.

Stimulate birth canal: The companion mouse interacts with the birth canal and external genital region of the dam in labor. This behavior includes direct contact by the companion mouse with the pregnant animal’s perineal region with the mouth or tongue or lateral pulling of the surrounding tissue using its forepaws.

Social interactions include:

Grooming: The companion mouse uses tongue, teeth, and forepaws to make repeated contact with the pregnant animal’s fur or skin. Grooming behaviors may target the other animal’s dorsal, lateral, or ventral surfaces.

Mounting: The companion mouse assumes a dorsal position on top of the pregnant female in labor and places its forepaws and body weight on the back to engage in rhythmic pelvic thrusting.

Some other behaviors (e.g., shepherding: companion mouse chases pregnant female to nest, encouraging the female in labor to enter and remain in the nest) were scored but were in practice rare and had a negligible contribution to outcomes across companion types, thus were not included in the transition matrices.

Inattentive behaviors include prolonged periods of disengagement, inactivity, or other self-aimed behaviors such as foraging, eating, self-grooming, sleeping, etc. To more accurately represent the temporal dynamics of assistive behaviors, we defined and classified inattention using the median duration of bouts where the companion animal was not interacting with the parturient female as a threshold, such that periods shorter than or equal to this cutoff were considered brief pauses in engagement, while those exceeding it were catalogued as true inattentiveness. The companion animals, especially the lactating wild-types, engaged in complex, energetically demanding, adaptive actions. This method accounted for natural breaks that occurred as part of the labor assistance process while making sure that substantial interruptions in social or prosocial behaviors were properly identified. We wanted to ensure this classification was informed by an empirical distribution instead of an arbitrary boundary. After behavioral classification, we calculated the transition matrix for the midwife behavior for the full duration of each of the births. We considered each behavior a discrete state, and transitions were computed to represent the likelihood of moving from one state to another. In this case we included self-loops in the transition diagrams to emphasize the potential importance of the behavioral pauses for labor assistance.

## Supplemental Information

**Figures S1** (related to Figure 2) and **S2** (related to Figure 4).

**Tables S1** and **S2** (related to Figure 2E), S3 and S4 (related to Figure 3C), **S5** and **S6** (related to Figure 3F), **S7** (related to Figure 4G), **S8** (related to Figure 4H), and **S9** (related to Figure 4I).

**Videos S1** (related to Figure 1A), **S2** (related to Figure 2B), **S3** (related to Figure 3), and **S4** and **S5** (related to Figure 4).

Source data spreadsheet containing data and statistical analyses for all figures, supplementary figures, and supplementary tables.

---

**Figure S1.**
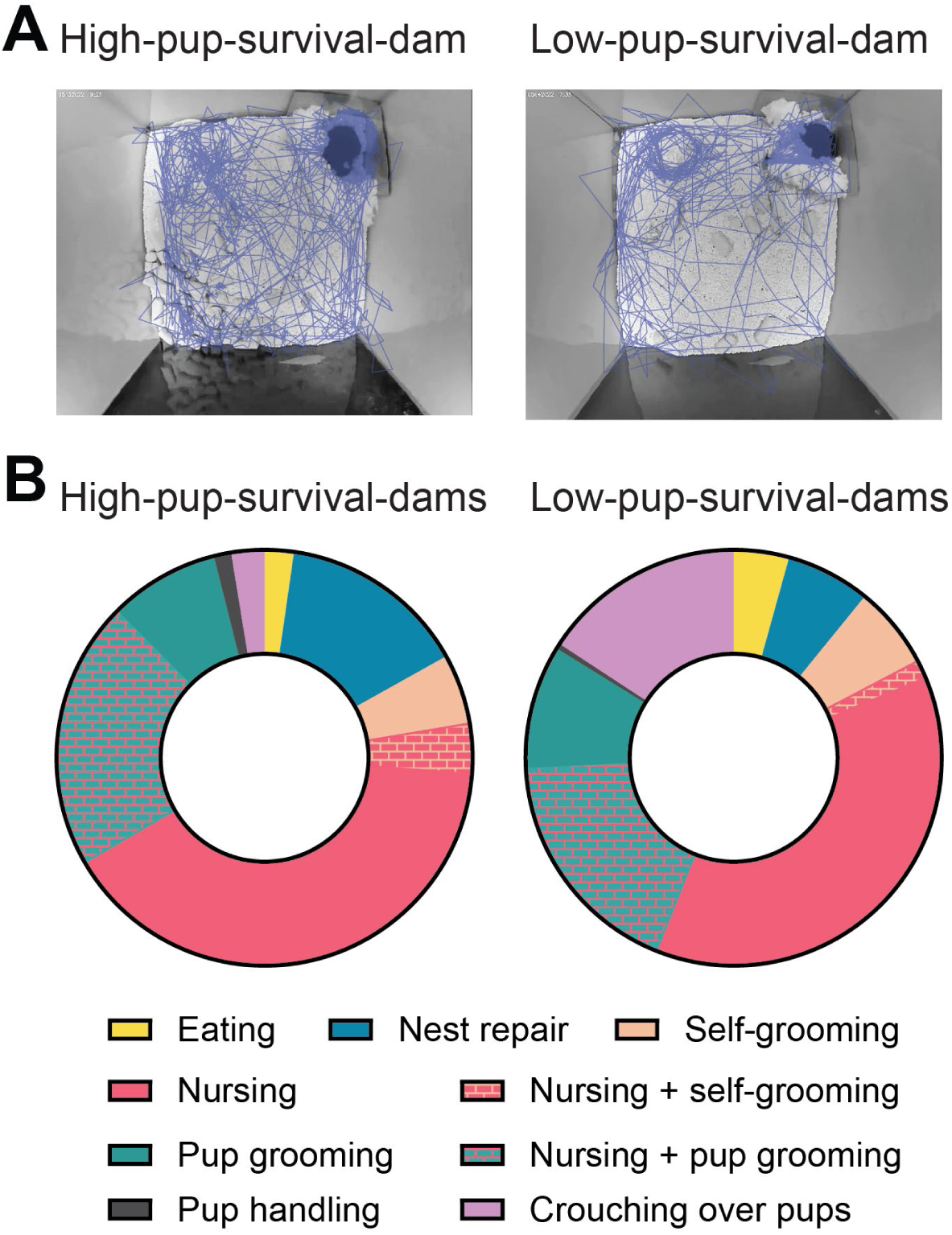
Analysis of time out-of-nest and in-nest behaviors. Related to. **Figure 2**. (A) Example of automated motion tracking for low-pup-survival dam to quantify time in nest and out of nest (sampled every second based on pose-estimate centroid). Blue lines, dam trajectory over 3 hours overlaid on video frame of enclosure. Dark gray, average nest location. Enclosure dimensions: 28×28×28 cm^3^. Left, high-pup-survival dam (total distance traveled: 7.85 meters; time spent in nest: 152.4 minutes or ∼85% of total time; time spent out of nest: 27.6 minutes or ∼15% total time). Right, low-pup-survival dam (total distance traveled: 35.75 meters; time spent in nest: 167.5 minutes or ∼93% of total time; time spent out of nest: 12.5 minutes or ∼7% of total time). (B) Time spent for in-nest behaviors for N=4 high-pup-survival and N=4 low-pup-survival dams.

**Figure S2.**
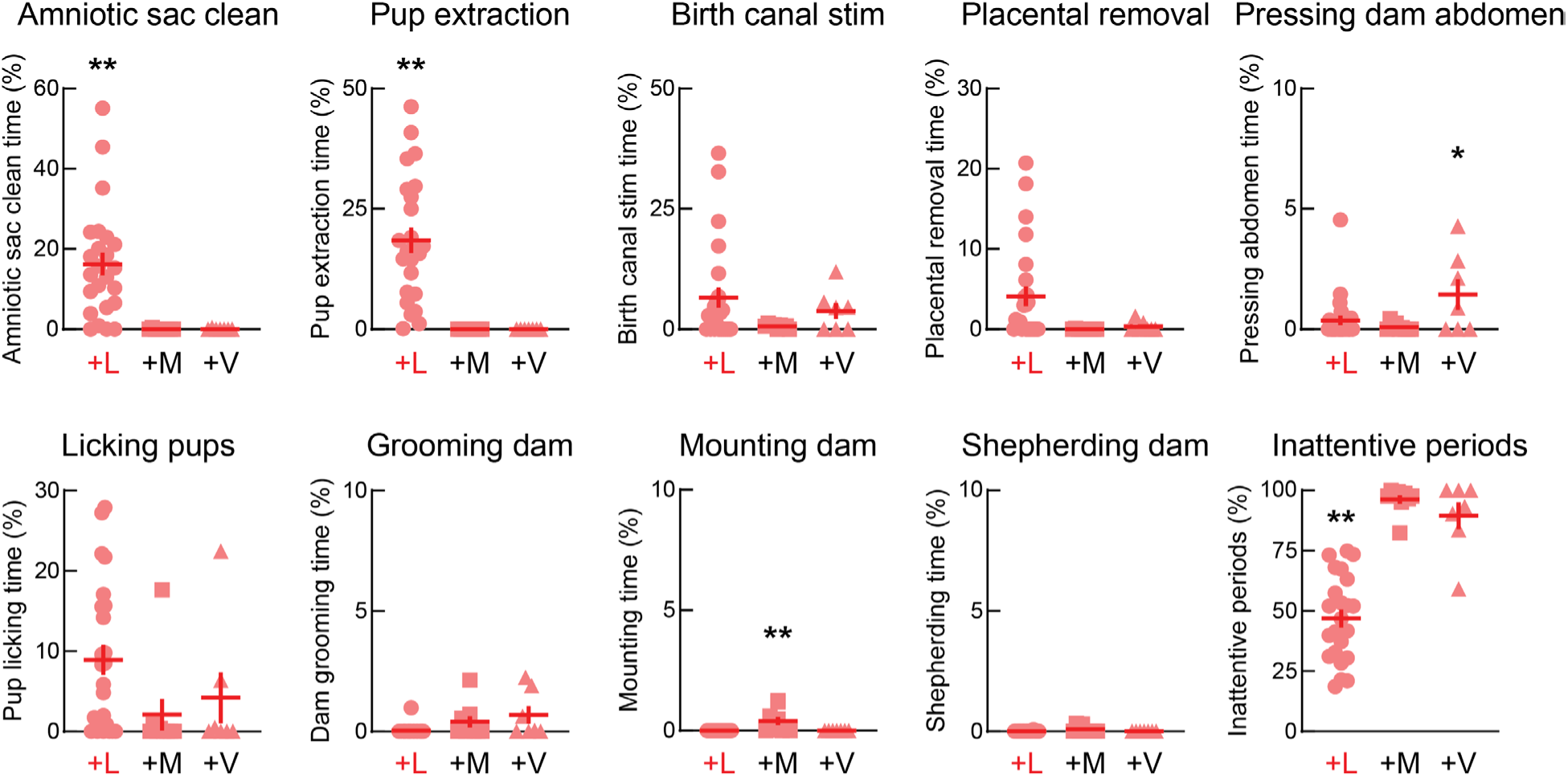
Relative time for behaviors of companion animal interacting with pregnant animal. Related to Figure 4. Each symbol represents relative fraction of time spent engaging in a given behavior averaged across each birthing of individual pups. ‘+L’, companion lactating female wild-type (N=5 dams, 23 pups); ‘+M’, companion paternal male wild-type (N=3 dams, 9 pups); ‘+V’, companion virgin female wild-type (N=3 dams, 7 pups). Lactating female companions spent significantly more time cleaning pups of amniotic sac (16.2±2.8%, p<0.005, ANOVA with Tukey’s multiple comparisons correction) than paternal males (0.5±0.5%) or virgin females (0.5±0.5%), spent more time extracting pups (+L: 18.4±2.6%, +M: 0%, +V: 0%, p<0.0005), and spent less time inattentive (+L: 46.9±3.7%, +M: 96.3±1.8%, +V: 89.5±5.6%, p<0.0001). Males spent more time mounting than females (+L: 0%, +M: 0.4±0.2%, +V: 0%, p<0.006). Virgin females spent more time pressing the abdomen than lactating females or males (+L: 0.4±0.2%, +M: 0.9±0.5%, +V: 1.4±0.6%, p<0.05).

**Supplementary Table S1.** High-pup-survival dam single parenting behavioral transitions. Related to Figure 2E.

| To \ From | In nest | Nest repair | Foraging | Pup check-ins | Nest buffering | Out of nest |
| --- | --- | --- | --- | --- | --- | --- |
| In nest |  | 0.80±0.03 | 0.06±0.02 | 0.18±0.04 | 0.13±0.04 | 0.20±0.03 |
| Nest repair | 0.68±0.03 |  | 0.03±0.02 | 0.18±0.04 | 0.17±0.04 | 0.06±0.01 |
| Foraging | 0.02±0.01 | 0.02±0.01 |  | 0.01±0.01 | 0.07±0.03 | 0.33±0.03 |
| Pup check-ins | 0.0±0.0 | 0.02±0.01 | 0.03±0.01 |  | 0.04±0.03 | 0.34±0.05 |
| Nest buffering | 0.04±0.01 | 0.03±0.01 | 0.02±0.01 | 0.02±0.01 |  | 0.07±0.01 |
| Out of nest | 0.26±0.03 | 0.13±0.02 | 0.86±0.03 | 0.61±0.05 | 0.59±0.08 |  |

**Supplementary Table S2.** Low-pup-survival dam single parenting behavioral transitions. Related to Figure 2E.

| To \ From | In nest | Nest repair | Foraging | Pup check-ins | Nest buffering | Out of nest |
| --- | --- | --- | --- | --- | --- | --- |
| In nest |  | 0.71±0.04 | 0.27±0.15 | 0.02±0.01 | 0.19±0.09 | 0.25±0.03 |
| Nest repair | 0.63±0.03 |  | 0.01±0.01 | 0.05±0.03 | 0.05±0.05 | 0.14±0.05 |
| Foraging | 0.06±0.04 | 0.02±0.01 |  | 0.0±0.0 | 0.0±0.0 | 0.27±0.06 |
| Pup check-ins | 0.0±0.0 | 0.02±0.01 | 0.0±0.0 |  | 0.0±0.0 | 0.28±0.08 |
| Nest buffering | 0.02±0.01 | 0.01±0.01 | 0.0±0.0 | 0.0±0.0 |  | 0.06±0.02 |
| Out of nest | 0.29±0.01 | 0.24±0.03 | 0.72±0.15 | 0.93±0.03 | 0.76±0.08 |  |

**Supplementary Table S3.** Initially low-pup-survival dam single parenting behavioral transitions for litter 2 before co-housing. Related to Figure 3C.

| From<br>To | In nest | Nest<br>repair | Foraging | Pup<br>check-ins | Nest<br>buffering | Out of nest |
| --- | --- | --- | --- | --- | --- | --- |
| In nest |  | 0.81±0.06 | 0.16±0.10 | 0.08±0.03 | 0.23±0.08 | 0.21±0.04 |
| Nest repair | 0.63±0.03 |  | 0.01±0.01 | 0.06±0.02 | 0.16±0.07 | 0.09±0.06 |
| Foraging | 0.03±0.02 | 0.01±0.00 |  | 0.01±0.01 | 0.02±0.01 | 0.40±0.07 |
| Pup check-ins | 0.0±0.0 | 0.0±0.0 | 0.02±0.01 |  | 0.02±0.01 | 0.27±0.04 |
| Nest buffering | 0.04±0.01 | 0.01±0.01 | 0.00±0.0 | 0.0±0.0 |  | 0.03±0.01 |
| Out of nest | 0.30±0.04 | 0.17±0.06 | 0.81±0.09 | 0.85±0.04 | 0.57±0.12 |  |

**Supplementary Table S4.** Initially low-pup-survival dam single parenting behavioral transitions for litter 3 after co-housing with high-pup-survival dam and litter. Related to Figure 3C.

| From<br>To | In nest | Nest<br>repair | Foraging | Pup<br>check-ins | Nest<br>buffering | Out of nest |
| --- | --- | --- | --- | --- | --- | --- |
| In nest |  | 0.85±0.04 | 0.10±0.09 | 0.33±0.08 | 0.19±0.08 | 0.22±0.04 |
| Nest repair | 0.66±0.03 |  | 0.0±0.01 | 0.20±0.05 | 0.08±0.03 | 0.02±0.01 |
| Foraging | 0.01±0.01 | 0.01±0.01 |  | 0.0±0.0 | 0.09±0.03 | 0.38±0.08 |
| Pup check-ins | 0.01±0.0 | 0.02±0.01 | 0.0±0.01 |  | 0.01±0.01 | 0.32±0.06 |
| Nest buffering | 0.02±0.01 | 0.02±0.01 | 0.0±0.0 | 0.01±0.01 |  | 0.06±0.03 |
| Out of nest | 0.30±0.04 | 0.10±0.04 | 0.90±0.09 | 0.46±0.11 | 0.63±0.16 |  |

**Supplementary Table S5.** Initially low-pup-survival dam single parenting behavioral transitions for litter 3 after co-housing with paternal male. Related to Figure 3F.

| To \ From | In nest | Nest repair | Foraging | Pup check-ins | Nest buffering | Out of nest |
| --- | --- | --- | --- | --- | --- | --- |
| In nest |  | 0.83±0.11 | 0.31±0.24 | 0.03±0.02 | 0.17±0.12 | 0.25±0.07 |
| Nest repair | 0.64±0.06 |  | 0.0±0.0 | 0.13±0.06 | 0.0±0.0 | 0.05±0.02 |
| Foraging | 0.12±0.10 | 0.04±0.04 |  | 0.0±0.0 | 0.0±0.0 | 0.35±0.14 |
| Pup check-ins | 0.0±0.0 | 0.0±0.0 | 0.0±0.0 |  | 0.0±0.0 | 0.32±0.11 |
| Nest buffering | 0.0±0.0 | 0.0±0.0 | 0.06±0.06 | 0.0±0.0 |  | 0.03±0.01 |
| Out of nest | 0.24±0.10 | 0.13±0.12 | 0.63±0.24 | 0.84±0.22 | 0.83±0.24 |  |

**Supplementary Table S6.**
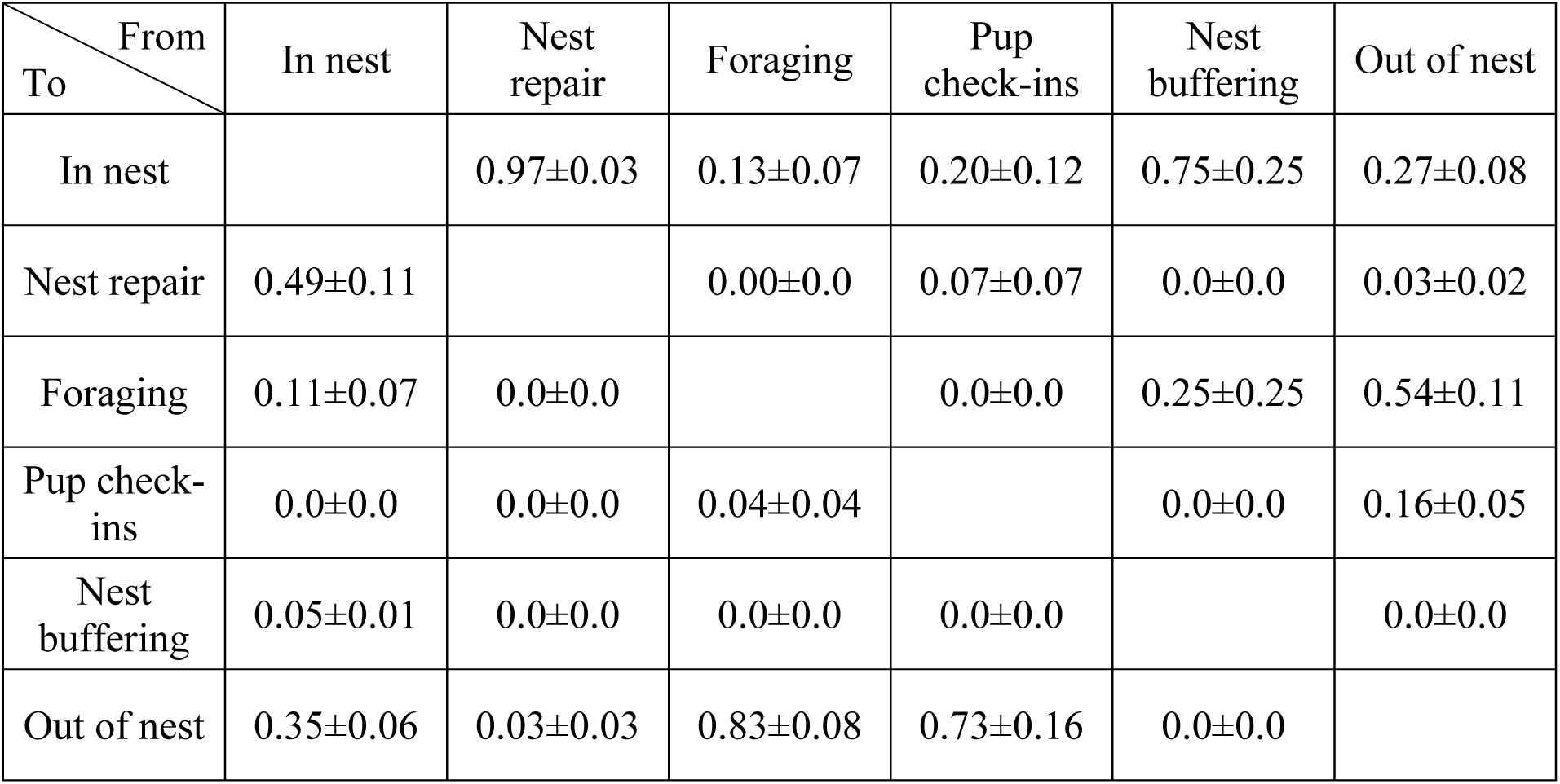
Initially low-pup-survival dam single parenting behavioral transitions for litter 3 after co-housing with nulliparous female. Related to Figure 3F.

**Supplementary Table S7.** Companion lactating wild-type female interactions with pregnant oxytocin receptor knockout dam. Related to Figure 4G.

| From<br>To | Placenta<br>removal | Amniotic<br>sac<br>clean | Stim<br>birth<br>canal | Groom | Inattent | Mount | Press<br>abdomen | Extract<br>pup | Lick<br>pup |
| --- | --- | --- | --- | --- | --- | --- | --- | --- | --- |
| Placenta<br>removal | 0.04±<br>0.04 | 0.04±<br>0.04 | 0.14±<br>0.07 | 0.0±<br>0.0 | 0.03±<br>0.02 | 0.0±<br>0.0 | 0.0±<br>0.0 | 0.15±<br>0.06 | 0.01±<br>0.01 |
| Amniotic<br>sac<br>clean | 0.0±<br>0.0 | 0.16±<br>0.04 | 0.10±<br>0.05 | 0.0±<br>0.0 | 0.24±<br>0.07 | 0.0±<br>0.0 | 0.08±<br>0.05 | 0.32±<br>0.07 | 0.03±<br>0.02 |
| Stim<br>birth<br>canal | 0.0±<br>0.0 | 0.09±<br>0.05 | 0.06±<br>0.03 | 0.02±<br>0.02 | 0.09±<br>0.03 | 0.0±<br>0.0 | 0.05±<br>0.03 | 0.08±<br>0.05 | 0.02±<br>0.01 |
| Groom | 0.0±<br>0.0 | 0.0±<br>0.0 | 0.01±<br>0.01 | 0.0±<br>0.0 | 0.01±<br>0.01 | 0.0±<br>0.0 | 0.0±<br>0.0 | 0.0±<br>0.0 | 0.0±<br>0.0 |
| Inattent | 0.0±<br>0.0 | 0.23±<br>0.06 | 0.24±<br>0.07 | 0.07±<br>0.05 | 0.0±<br>0.0 | 0.0±<br>0.0 | 0.10±<br>0.04 | 0.24±<br>0.06 | 0.36±<br>0.08 |
| Mount | 0.0±<br>0.0 | 0.0±<br>0.0 | 0.0±<br>0.0 | 0.0±<br>0.0 | 0.0±<br>0.0 | 0.0±<br>0.0 | 0.0±<br>0.0 | 0.0±<br>0.0 | 0.0±<br>0.0 |
| Press<br>abdomen | 0.0±<br>0.0 | 0.02±<br>0.01 | 0.01±<br>0.01 | 0.0±<br>0.0 | 0.03±<br>0.02 | 0.0±<br>0.0 | 0.04±<br>0.02 | 0.0±<br>0.0 | 0.01±<br>0.01 |
| Extract<br>pup | 0.0±<br>0.0 | 0.23±<br>0.06 | 0.01±<br>0.01 | 0.0±<br>0.0 | 0.33±<br>0.08 | 0.0±<br>0.0 | 0.01±<br>0.01 | 0.18±<br>0.05 | 0.14±<br>0.07 |
| Lick<br>pup | 0.0±<br>0.0 | 0.04±<br>0.02 | 0.05±<br>0.02 | 0.0±<br>0.0 | 0.23±<br>0.05 | 0.0±<br>0.0 | 0.02±<br>0.02 | 0.03±<br>0.02 | 0.17±<br>0.04 |

**Supplementary Table S8.**
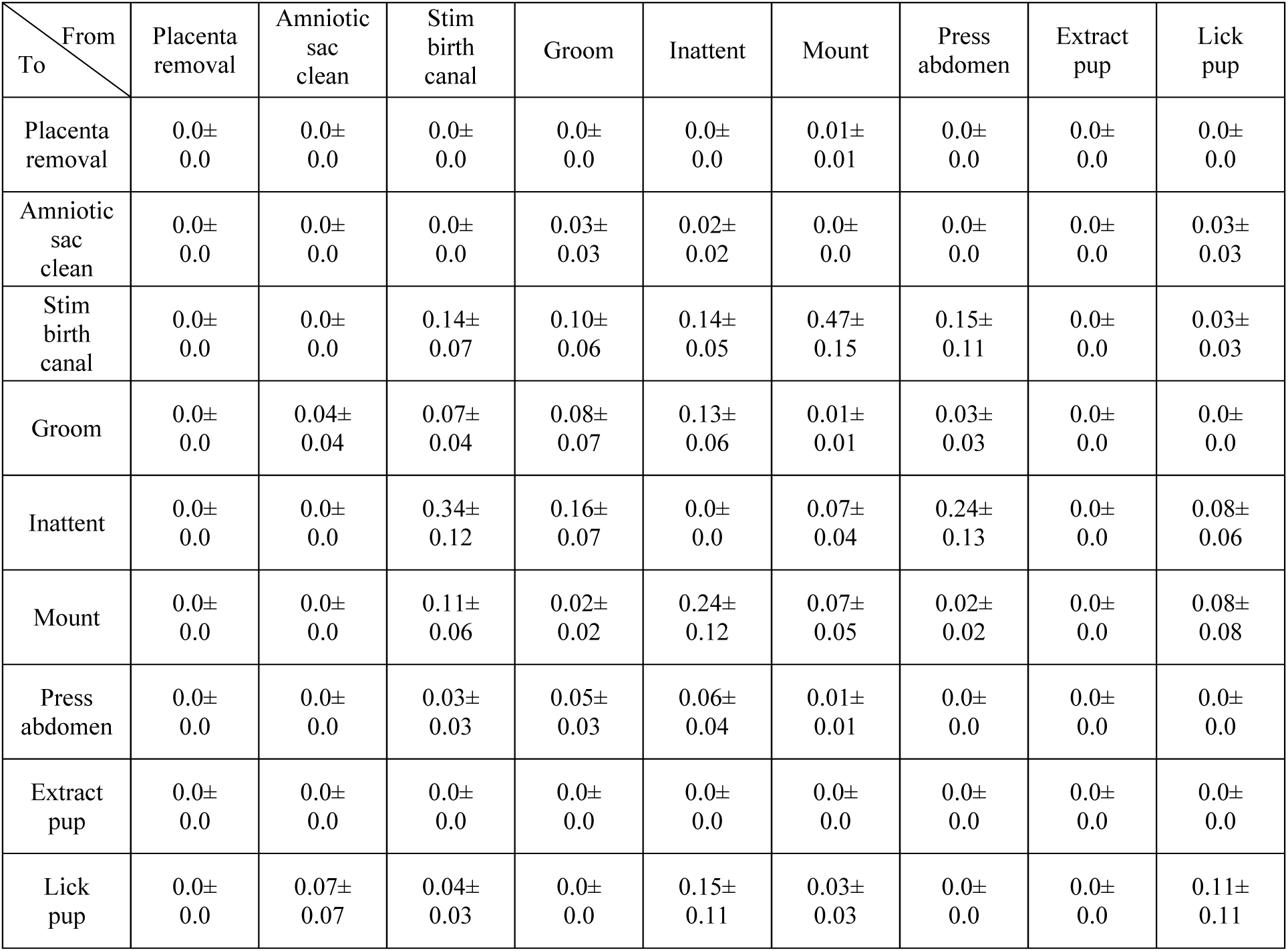
Companion wild-type paternal male interactions with pregnant oxytocin receptor knockout dam. Related to Figure 4H.

| From<br>To | Placenta<br>removal | Amniotic<br>sac<br>clean | Stim<br>birth<br>canal | Groom | Inattent | Mount | Press<br>abdomen | Extract<br>pup | Lick<br>pup |
| --- | --- | --- | --- | --- | --- | --- | --- | --- | --- |
| Placenta<br>removal | 0.0±<br>0.0 | 0.0±<br>0.0 | 0.0±<br>0.0 | 0.0±<br>0.0 | 0.0±<br>0.0 | 0.01±<br>0.01 | 0.0±<br>0.0 | 0.0±<br>0.0 | 0.0±<br>0.0 |
| Amniotic<br>sac<br>clean | 0.0±<br>0.0 | 0.0±<br>0.0 | 0.0±<br>0.0 | 0.03±<br>0.03 | 0.02±<br>0.02 | 0.0±<br>0.0 | 0.0±<br>0.0 | 0.0±<br>0.0 | 0.03±<br>0.03 |
| Stim<br>birth<br>canal | 0.0±<br>0.0 | 0.0±<br>0.0 | 0.14±<br>0.07 | 0.10±<br>0.06 | 0.14±<br>0.05 | 0.47±<br>0.15 | 0.15±<br>0.11 | 0.0±<br>0.0 | 0.03±<br>0.03 |
| Groom | 0.0±<br>0.0 | 0.04±<br>0.04 | 0.07±<br>0.04 | 0.08±<br>0.07 | 0.13±<br>0.06 | 0.01±<br>0.01 | 0.03±<br>0.03 | 0.0±<br>0.0 | 0.0±<br>0.0 |
| Inattent | 0.0±<br>0.0 | 0.0±<br>0.0 | 0.34±<br>0.12 | 0.16±<br>0.07 | 0.0±<br>0.0 | 0.07±<br>0.04 | 0.24±<br>0.13 | 0.0±<br>0.0 | 0.08±<br>0.06 |
| Mount | 0.0±<br>0.0 | 0.0±<br>0.0 | 0.11±<br>0.06 | 0.02±<br>0.02 | 0.24±<br>0.12 | 0.07±<br>0.05 | 0.02±<br>0.02 | 0.0±<br>0.0 | 0.08±<br>0.08 |
| Press<br>abdomen | 0.0±<br>0.0 | 0.0±<br>0.0 | 0.03±<br>0.03 | 0.05±<br>0.03 | 0.06±<br>0.04 | 0.01±<br>0.01 | 0.0±<br>0.0 | 0.0±<br>0.0 | 0.0±<br>0.0 |
| Extract<br>pup | 0.0±<br>0.0 | 0.0±<br>0.0 | 0.0±<br>0.0 | 0.0±<br>0.0 | 0.0±<br>0.0 | 0.0±<br>0.0 | 0.0±<br>0.0 | 0.0±<br>0.0 | 0.0±<br>0.0 |
| Lick<br>pup | 0.0±<br>0.0 | 0.07±<br>0.07 | 0.04±<br>0.03 | 0.0±<br>0.0 | 0.15±<br>0.11 | 0.03±<br>0.03 | 0.0±<br>0.0 | 0.0±<br>0.0 | 0.11±<br>0.11 |

**Supplementary Table S9.**
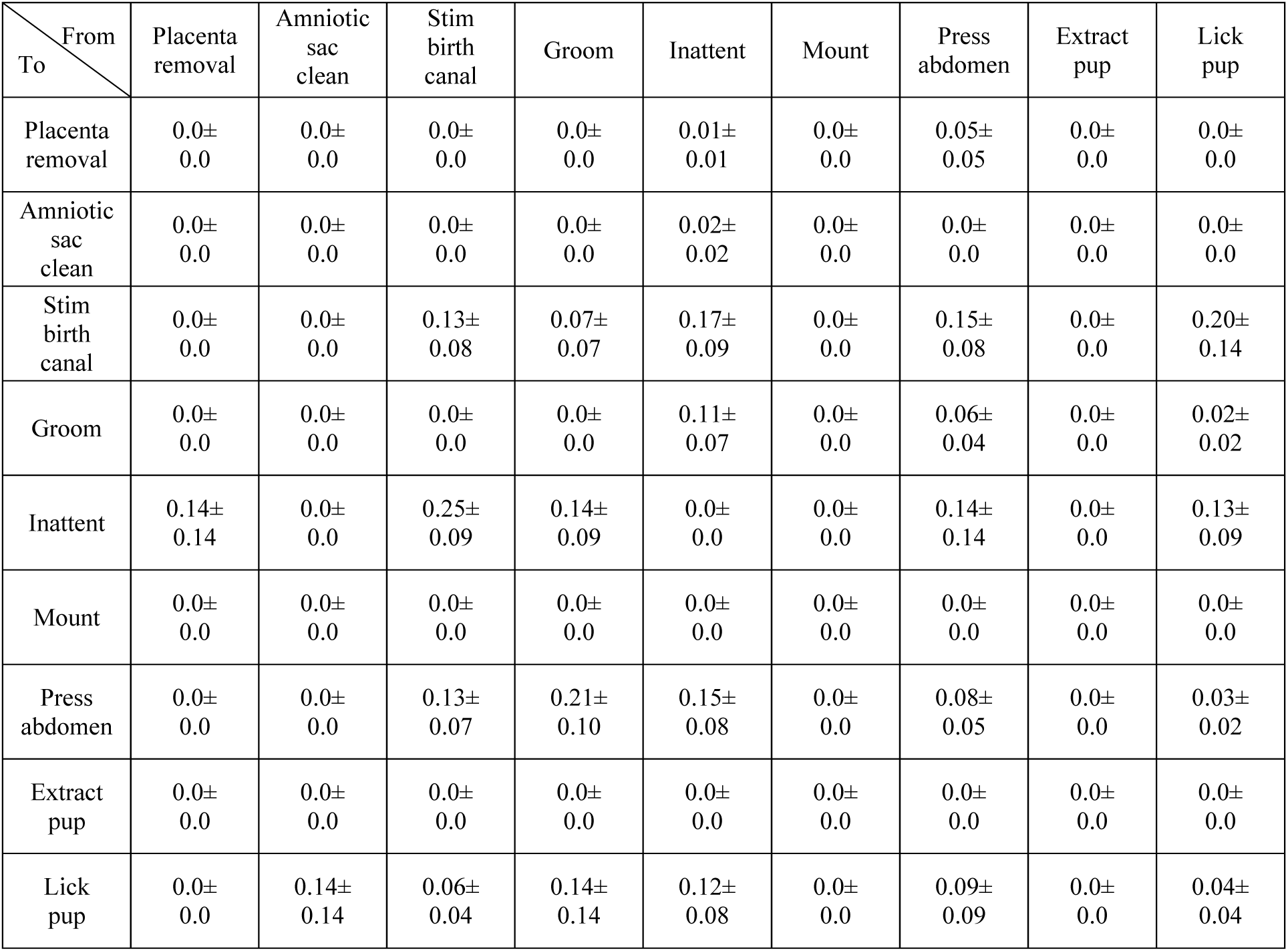
Companion nulliparous wild-type female interactions with pregnant oxytocin receptor knockout dam. Related to Figure 4I.

| From<br>To | Placenta<br>removal | Amniotic<br>sac<br>clean | Stim<br>birth<br>canal | Groom | Inattent | Mount | Press<br>abdomen | Extract<br>pup | Lick<br>pup |
| --- | --- | --- | --- | --- | --- | --- | --- | --- | --- |
| Placenta<br>removal | 0.0±<br>0.0 | 0.0±<br>0.0 | 0.0±<br>0.0 | 0.0±<br>0.0 | 0.01±<br>0.01 | 0.0±<br>0.0 | 0.05±<br>0.05 | 0.0±<br>0.0 | 0.0±<br>0.0 |
| Amniotic<br>sac<br>clean | 0.0±<br>0.0 | 0.0±<br>0.0 | 0.0±<br>0.0 | 0.0±<br>0.0 | 0.02±<br>0.02 | 0.0±<br>0.0 | 0.0±<br>0.0 | 0.0±<br>0.0 | 0.0±<br>0.0 |
| Stim<br>birth<br>canal | 0.0±<br>0.0 | 0.0±<br>0.0 | 0.13±<br>0.08 | 0.07±<br>0.07 | 0.17±<br>0.09 | 0.0±<br>0.0 | 0.15±<br>0.08 | 0.0±<br>0.0 | 0.20±<br>0.14 |
| Groom | 0.0±<br>0.0 | 0.0±<br>0.0 | 0.0±<br>0.0 | 0.0±<br>0.0 | 0.11±<br>0.07 | 0.0±<br>0.0 | 0.06±<br>0.04 | 0.0±<br>0.0 | 0.02±<br>0.02 |
| Inattent | 0.14±<br>0.14 | 0.0±<br>0.0 | 0.25±<br>0.09 | 0.14±<br>0.09 | 0.0±<br>0.0 | 0.0±<br>0.0 | 0.14±<br>0.14 | 0.0±<br>0.0 | 0.13±<br>0.09 |
| Mount | 0.0±<br>0.0 | 0.0±<br>0.0 | 0.0±<br>0.0 | 0.0±<br>0.0 | 0.0±<br>0.0 | 0.0±<br>0.0 | 0.0±<br>0.0 | 0.0±<br>0.0 | 0.0±<br>0.0 |
| Press<br>abdomen | 0.0±<br>0.0 | 0.0±<br>0.0 | 0.13±<br>0.07 | 0.21±<br>0.10 | 0.15±<br>0.08 | 0.0±<br>0.0 | 0.08±<br>0.05 | 0.0±<br>0.0 | 0.03±<br>0.02 |
| Extract<br>pup | 0.0±<br>0.0 | 0.0±<br>0.0 | 0.0±<br>0.0 | 0.0±<br>0.0 | 0.0±<br>0.0 | 0.0±<br>0.0 | 0.0±<br>0.0 | 0.0±<br>0.0 | 0.0±<br>0.0 |
| Lick<br>pup | 0.0±<br>0.0 | 0.14±<br>0.14 | 0.06±<br>0.04 | 0.14±<br>0.14 | 0.12±<br>0.08 | 0.0±<br>0.0 | 0.09±<br>0.09 | 0.0±<br>0.0 | 0.04±<br>0.04 |

## Notes

### Competing Interest Statement

The authors have declared no competing interest.

### Summary of Updates

Major updates to data analysis and results.

## References

1. Dulac, C., O’Connell, L.A., & Wu, Z. (2014). Neural control of maternal and paternal behaviors. Science 345, 765–770.

2. Rilling, J.K., and Young, L.J. (2014). The biology of mammalian parenting and its effect on offspring social development. Science 345, 771–776.

3. Leon, M., Croskerry, P.G., and Smith, G.K. (1978). Thermal control of mother-young contact in rats. Physiol. Behav. 21, 793–811.

4. Lynch, C.B., and Possidente, B.P. (1978). Relationships of maternal nesting to thermoregulatory nesting in house mice (mus musculus) at warm and cold temperatures. Animal Behav. 26, 1136–1143.

5. Gaskill, B.N., Gordon, C.J., Pajor, E.A., Lucas, J.R., Davis, J.K., and Garner, J.P. (2012). Heat or insulation: Behavioral titration of mouse preference for warmth or access to a nest. PLoS ONE 7, e32799.

6. König, B. (1993). Maternal investment of communally nursing female house mice (Mus musculus domesticus). Behav. Processes 30, 61–73.

7. Manning, C.J., Dewsbury, D.A, Wakeland, E.K., and Potts, W.K. (1995) Communal nesting and communal nursing in house mice, Mus musculus domesticus. Animal Behav. 50, 741–751.

8. Ferrari, M., Lindholm, A.K., & König, B. (2019). Fitness consequences of female alternative reproductive tactics in house mice (*mus musculus domesticus*). Am. Nat. 193, 106–124.

9. Capas-Peneda, S., Morello, G.M., Lamas, S., Olsson, I.A.S., and Gilbert, C. (2020). Causes of death in newborn C57BL/6J mice. bioRxiv doi:10.1101/2020.02.25.964551.

10. Brajon, S., Morello, G.M., Capas-Peneda, S., Hultgren, J., Gilbert, C., and Olsson, A. (2021). All the pups we cannot see: Cannibalism masks perinatal death in laboratory mouse breeding but infanticide is rare. Animals 11, 2327.

11. Richard, P., Moss, F., and Freund-Mercier, M.J. (1991). Central effects of oxytocin. Physiol. Rev. 71, 331–370.

12. Gimpl, G., and Fahrenholz, F. (2001). The oxytocin receptor system: structure, function, and regulation. Physiol. Rev. 81, 629–683.

13. Jurek, B., & Neumann, I.D. (2018). The oxytocin receptor: from intracellular signaling to behavior. Physiol. Rev. 98, 1805–1908.

14. Theofanopoulou, C., Gedman, G., Cahill, J.A., Boeckx, C., and Jarvis, E.D. (2021). A universal evolution-based nomenclature for the oxytocin and vasotocin ligand and receptor families. Nature 592, 747–755.

15. Tang, Y., Benusiglio, D., Lefevre, A., Hilfiger, L., Althammer, F., Bludau, A., Hagiwara, D., Baudon, A., Darbon, P., Schimmer, J., Kirchner, M.K., Roy, R.K., Wang, S., Eliava, M., Wagner, S., Oberhuber, M., Conzelmann, K.K., Schwarz, M., Stern, J.E., Leng, G., Neumann, I.D., Charlet, A., & Grinevich, V. (2020). Social touch promotes interfemale communication via activation of parvocellular oxytocin neurons. Nat. Neurosci. 23, 1125–1137.

16. Carcea, I., Caraballo, N.L., Marlin, B.J., Ooyama, R., Riceberg, J.S., Mendoza Navarro, J.M., Opendak, M., Diaz, V.E., Schuster, L., Alvarado Torres, M.I., Lethin, H., Ramos, D., Minder, J., Mendoza, S.L., Bair-Marshall, C.J., Samadjopoulos, G.H., Hidema, S., Falkner, A., Lin, D., Mar, A., Wadghiri, Y.Z., Nishimori, K., Kikusui, T., Mogi, K., Sullivan, R.M., and Froemke, R.C. (2021). Oxytocin neurons enable social transmission of maternal behaviour. Nature 596, 553–557.

17. McNeilly, A.S., Robinson, I.C., Houston, M.J., and Howie, P.W. (1983) Release of oxytocin and prolactin in response to suckling. Br. Med. J. 286, 257–259.

18. Valtcheva, S., Issa, H.A., Bair-Marshall, C.J., Martin, K.A., Jung, K., Zhang, Y., Kwon, H.-B., and Froemke, R.C. (2023). Neural circuitry for maternal oxytocin release induced by infant cries. Nature 621, 788–795.

19. Froemke, R.C. and Young, L.J. Oxytocin modulation and neural plasticity. (2021) Annu. Rev. Neurosci. 44, 359–381.

20. Takayanagi, Y., Yoshida, M., Bielsky, I.F., Ross, H.E., Kawamata, M., Onaka, T., Yanagisawa, T., Kimura, T., Matzuk, M. M., Young, L. J., and Nishimori, K. (2005). Pervasive social deficits, but normal parturition, in oxytocin receptor-deficient mice. Proc. Natl. Acad. Sci. U S A 102:16096–16101.

21. Berendzen, K.M., Sharma, R., Mandujano, M.A., Wei, Y., Rogers, F.D., Simmons, T.C., Seelke, A.M.H., Bond, J.M., Larios, R., Goodwin, N.L., Sherman, M., Parthasarthy, S., Espineda, I., Knoedler, J.R., Beery, A., Bales, K.L., Shah, N.M., and Manoli, D.S. (2023). Oxytocin receptor is not required for social attachment in prairie voles. Neuron 111, 787–796.

22. Weber, E.M., Algers, B., Hultgren, J., and Olsson, I.A.S. (2013). Pup mortality in laboratory mice—Infanticide or not? Acta Veterinaria Scandinavica 55, 83.

23. Bronikowski, A.M., Cords, M., Alberts, S.C., Altmann, J., Brockman, D.K., Fedigan, L.M., Pusey, A., Stoinski, T., Strier, K.B., and Morris, W.F. (2016). Female and male life tables for seven wild primate species. Scientific Data 3, 160006.

24. Dwyer, C.M., Conington, J., Corbiere, F., Holmøy, I.H., Muri, K., Nowak, R., Rooke, J., Vipond, J., & Gautier, J.-M. (2016). Improving neonatal survival in small ruminants: science into practice. Animal 10, 449–459.

25. 25. Heldstab, S.A., van Schaik, C.P., Müller, D.W.H., Rensch, E., Lackey, L.B., Zerbe, P., Hatt, J.-M., Clauss, M., and Matsuda, I. (2021). Reproductive seasonality in primates: Patterns, concepts and unsolved questions. Biol. Rev. 96, 66–88.

26. Perani, C.V., and Slattery, D.A. (2014). Using animal models to study post-partum psychiatric disorders. Br. J. Pharmacol. 171, 4539–4555.

27. Mir, F.R., Pollano, A., and Rivarola, M.A. (2022). Animal models of postpartum depression revisited. Psychoneuroendocrinol. 136, 105590.

28. Król E., and Speakman, J.R. (2003). Limits to sustained energy intake VI. energetics of lactation in laboratory mice at thermoneutrality. J. Exp. Biol. 206, 4255–4266.

29. Gamo, Y., Bernard, A., Troup, C., Munro, F., Derrer, K., Jeannesson, N., Campbell, A., Gray, H., Miller, J., Dixon, J., Mitchell, S.E., Hambly, C., Vaanholt, L.M., and Speakman, J.R. (2016). Limits to sustained energy intake XXIV: Impact of suckling behaviour on the body temperatures of lactating female mice. Sci. Rep. 6, 25665.

30. Król, E., and Speakman, J.R. (2019). Switching off the furnace: brown adipose tissue and lactation. Mol. Aspects Med. 68, 18–41.

31. Marlin, B.J., Mitre, M., D’amour, J.A., Chao, M.V., and Froemke, R.C. (2015). Oxytocin enables maternal behaviour by balancing cortical inhibition. Nature 520, 499–504.

32. Bussell, J.J., and Vosshall, L.B. (2012). Learning to suckle with signature odor. Curr. Biol. 22, 907–909.

33. Bernard, K., Simons, R., and Dozier, M. (2015). Effects of an attachment-based intervention on child protective services-referred mothers’ event-related potentials to children’s emotions. Child Dev. 86, 1673–1684.

34. Cohen, N., and Katz, C. (2021). Preventing child maltreatment: Key conclusions from a systematic literature review of prevention programs for practitioners. Child Abuse Negl. 118, 105138.

35. U.S. Department of Health and Human Services, Administration for Children and Families, Administration on Children, Youth and Families, Children’s Bureau. (2024). Child Maltreatment 2023.

36. Sobczak, A., Taylor, L., Solomon, S., Ho, J., Kemper, S., Phillips, B., Jacobson, K., Castellano, C., Ring, A., Castellano, B., and Jacobs, R.J. (2023). The effect of doulas on maternal and birth outcomes: a scoping review. Cureus 15, e39451.

37. Langford, D.J., Crager, S.E., Shehzad, Z., Smith, S.B., Sotocinal, S.G., Levenstadt, J.S., Chanda, M.L., Levitin, D.J., and Mogil, J.S. (2006). Social modulation of pain as evidence for empathy in mice. Science 312, 1967–1970.

38. Ben-Ami Bartal, I., Shan, H., Molasky, N.M.R., Murray, T.M., Williams, J.Z., Decety, J., and Mason, P. (2016). Anxiolytic treatment impairs helping behavior in rats. Front Psychol. 7, 850.

39. Burkett, J.P., Andari, E., Johnson, Z.V., Curry, D.C., de Waal, F.B.M., and Young, L.J. (2016). Oxytocin-dependent consolation behavior in rodents. Science 351, 375–378.

40. Carneiro de Oliveira, P.E., Carmona, I.M., Casarotto, M., Silveira, L.M., Oliveira, A.C.B., and Canto-de-Souza, A. (2022). Mice cohabiting with familiar conspecific in chronic stress condition exhibit methamphetamine-induced locomotor sensitization and augmented consolation behavior. Front. Behav. Neurosci. 16, 835717.

41. Zhang, M., Wu, Y.E., Jiang, M., and Hong, W. (2024). Cortical regulation of helping behaviour towards others in pain. Nature 626, 136–144.

42. Cao, P., Liu, Y., Ni, Z., Zhang, M., Wei, H.R., Liu, A., Guo, J.R., Yang, Y., Xu, Z., Guo, Y., Zhang, Z., Tao, W., & Wang, L. (2025). Rescue-like behavior in a bystander mouse toward anesthetized conspecifics promotes arousal via a tongue-brain connection. Sci. Adv. 11, eadq3874.

43. Sun, F., Wu, Y.E., & Hong, W. (2025). A neural basis for prosocial behavior toward unresponsive individuals. Science 387, eadq2679.

44. Sun, W., Zhang, G.W., Huang, J.J., Tao, C., Seo, M.B., Tao, H.W., & Zhang, L.I. (2025). Reviving-like prosocial behavior in response to unconscious or dead conspecifics in rodents. Science 387, eadq2677.

45. Ward, R. (1980). Some effects of strain differences in the maternal behavior of inbred mice. Devel. Psychobiol. 13, 181–190.

46. Bendesky, A., Kwon, Y.M., Lassance, J.M., Lewarch, C.L., Yao, S., Peterson, B.K., He, M.X., Dulac, C., & Hoekstra, H.E. (2017). The genetic basis of parental care evolution in monogamous mice. Nature 544, 434–439.

47. Fleszar, L.G., Bryant, A.S., Johnson, C.O., Blacker, B.F., Aravkin, A., Baumann, M., Dwyer-Lindgren, L., Kelly, Y.O., Maass, K., Zheng, P., and Roth, G.A. (2023). Trends in state-level maternal mortality by racial and ethnic group in the United States. JAMA 330, 52–61.

48. Volk, A.A., and Atkinson, J.A. (2013). Infant and child death in the human environment of evolutionary adaptation. Evol. Human Behav. 34, 182–192.

49. Benedict, J., and Cudmore, R.H. (2023). PiE: an open-source pipeline for home cage behavioral analysis. Front. Neurosci. 17, 1222644.

50. Mathis, A., Mamidanna, P., Cury, K.M., Abe, T., Murthy, V.N., Mathis, M.W., and Bethge, M. (2018). DeepLabCut: markerless pose estimation of user-defined body parts with deep learning. Nat. Neurosci. 21, 1281–1289.

51. Nath, T., Mathis, A., Chen, A.C., Patel, A., Bethge, M., and Mathis, M.W. (2019). Using DeepLabCut for 3D markerless pose estimation across species and behaviors. Nat. Protoc. 14, 2152–2176.

52. Datavyu Team (2014). Datavyu: A video coding tool. Databrary Project, New York University. URL: http://datavyu.org.

53. McHugh, M.L. (2012). Interrater reliability: the kappa statistic. Biochem. Med. 22, 276–282.

54. Pang, S.C., Janzen-Pang, J., Tse, M.Y., Croy, B.A., and Lima, P.D.A. (2014). The Cycling and Pregnant Mouse: Gross Anatomy. In B.A. Croy, A.T. Yamada, F.J. DeMayo, and S.L. Adamson (Eds.), The Guide to Investigation of Mouse Pregnancy (pp. 3–19). Academic Press.

55. Mota-Rojas, D., Orihuela, A., Strappini, A., Villanueva-García, D., Napolitano, F., Mora-Medina, P., Barrios-García, H.B., Herrera, Y., Lavalle, E., and Martínez-Burnes, J. (2020). Consumption of maternal placenta in humans and nonhuman mammals: beneficial and adverse effects. Animals 10, 2398.

